# Glia phagocytose neuronal sphingolipids to infiltrate developing synapses

**DOI:** 10.1101/2025.04.14.648777

**Authors:** Emma K. Theisen, Irma Magaly Rivas-Serna, Ryan J. Lee, Taylor R. Jay, Govind Kunduri, Tasha T. Nguyen, Vera Mazurak, M. Thomas Clandinin, Thomas R. Clandinin, John P. Vaughen

## Abstract

The complex morphologies of mature neurons and glia emerge through profound rearrangements of cell membranes during development. Despite being integral components of these membranes, it is unclear whether lipids might actively sculpt these morphogenic processes. By analyzing lipid levels in the developing fruit fly brain, we discover dramatic increases in specific sphingolipids coinciding with neural circuit establishment. Disrupting this sphingolipid bolus via genetic perturbations of sphingolipid biosynthesis and catabolism leads to impaired glial autophagy. Remarkably, glia can obtain sphingolipid precursors needed for autophagy by phagocytosing neurons. These precursors are then converted into specific long-chain ceramide phosphoethanolamines (CPEs), invertebrate analogs of sphingomyelin. These lipids are essential for glia to arborize and infiltrate the brain, a critical step in circuit maturation that when disrupted leads to reduced synapse numbers. Taken together, our results demonstrate how spatiotemporal tuning of sphingolipid metabolism during development plays an instructive role in programming brain architecture.

**Highlights:**

- Brain sphingolipids (SLs) remodel to very long-chain species during circuit maturation
- Glial autophagy requires *de novo* SL biosynthesis coordinated across neurons and glia
- Glia evade a biosynthetic blockade by phagolysosomal salvage of neuronal SLs
- Ceramide Phosphoethanolamine is critical for glial infiltration and synapse density

## Introduction

The dynamic regulation of cellular membranes in the brain is essential for circuit formation, synaptic function, and structural plasticity. During development, neurons extend and retract arbors, adding and removing membranes as they seek precise connections with synaptic partners^1–4^. In parallel, glial cells differentiate into diverse subtypes with distinctive and complex morphologies^5–9^. Previous work has described many proteins that regulate synaptic specificity and the establishment of glial architecture^10–17^. However, the possible roles of membrane lipids in shaping these processes remain largely unexplored.

Brain development requires the morphological maturation and precise integration of glial cells. In the mammalian brain, astrocytes extend thousands of fine processes throughout the neuropil, closely associating with synapses to modulate circuit function^7,18–20^. Oligodendrocytes wrap neuronal arbors to facilitate electrical conduction^21^, while microglia sculpt neural circuits through synaptic pruning^22^. In the *Drosophila* mushroom body, glial cells infiltrate just prior to engulfing fragmented axons, which are subsequently processed via the endolysosomal pathway^23^. Glial entry into the brain during early development and glial remodeling in response to axonal injury has been linked to phagocytosis^24,25^. However, the role of the lipidome in shaping glial arborization and infiltration is unknown.

Endolysosomal processing depends on lipid membranes to form vesicular structures that transition through a continuum of intermediates, culminating in lysosomes, the terminal degradation hubs of the cell. These transitions are marked by changes in membrane-associated proteins, such as the replacement of Rab5 by Rab7 during the transition from early endosomes to late endosomes^26^, as well as changes in lipid composition that together reflect endosomal identity. The addition of membrane material is crucial to the growing autophagosome, and lipid components of endosomal membranes can play signaling roles^21,27,28^. Finally, lipids are largely processed in the lysosome, where resident enzymes catabolize specific lipid classes^29–31^.

Brain lipids are highly diverse, with thousands of species varying within and across tissues^32–37^. The majority of a cell’s lipidome is comprised of glycerophospholipids including phosphatidylcholine, phosphoethanolamine, and phosphatidylserine, as well as sterols like cholesterol^38,39^. However, several rarer lipid classes exist that together account for less than 10% of brain lipids^34^. Intriguingly, mutations in enzymes that synthesize and catabolize one of these classes, the sphingolipids, are associated with a number of neurodegenerative disorders including Parkinson’s Disease, amyotrophic lateral sclerosis, and frontotemporal dementia^40–46^. Moreover, recent work in other systems has tied changes in sphingolipids to a wide variety of membrane rearrangements, including Purkinje and hippocampal neuron growth^47,48^, myelin wrapping and compaction^49–52^, and changes in microglia morphology^53,54^. In the fly brain, sphingolipids regulate myriad functions, spanning neuropil compartmentalization^55^ and neural circuit remodeling^56^ to rhodopsin trafficking^57,58^, neural excitability^59^, peripheral glial wrapping^60,61^, immune activation^62^, protein clearance^63–68^, and the control of circadian rhythms^65,69^. Some of these phenotypes have been associated with defects in endolysosomal processing. For example, *gba1b* mutants have enlarged lysosomes and accumulate the autophagy adaptor p62^63–65,70,71^. Previously, we found that sphingolipids directly program ultrastructural remodeling in circadian circuits, with low sphingolipids permitting neurite outgrowth, and high sphingolipids causing neurite retraction^65^. These observations raised the exciting hypothesis that dynamic changes in sphingolipid levels could be used to control membrane rearrangements.

To test this hypothesis, we sought to characterize sphingolipids during brain development, when many cells undergo dramatic morphological changes. Strikingly, we find that sphingolipid metabolism is dynamic, with a temporally restricted bolus of sphingolipids that emerges as synapses begin to assemble, and which produces long-chain sphingolipids as circuits mature and glia infiltrate. Disrupting sphingolipid catabolism specifically in glia impairs endolysosomal processing, while blocking *de novo* sphingolipid biosynthesis in both neurons and glia disrupts autophagy in glia. Strikingly, glia can acquire the sphingolipids they need for autophagy by phagocytosing neuronal membranes. When glia cannot synthesize a specific sphingolipid, Ceramide Phosphoethanolamine (CPE), they fail to infiltrate the brain, leading to reductions in synapse numbers. Thus, by the timely production and incorporation of neuronal sphingolipids, glia acquire the capacity to complete autophagy and to arborize into mature circuits.

## Results

### Sphingolipids are extensively remodeled during brain development

Given that sphingolipids play an instructive role in shaping neurite remodeling in circadian circuits in the adult brain, we hypothesized that sphingolipids might also sculpt earlier stages of brain development. In *Drosophila*, the adult brain begins to assemble during the third instar larval stage, with the first part of pupal development (0-24 hours after puparium formation (APF)) characterized by proliferation and differentiation of adult neurons and glia, as well as intensive glial phagocytosis of degenerating larval neuropils^23,24^ (Figure 1A). Next, differentiated neurites from adult neurons extend towards specific synaptic targets, beginning synapse assembly at approximately 48h APF. Over the second half of pupal development, synapse assembly, maturation and refinement continue^72–74^. Finally, in parallel with these later stages of neuronal maturation, specific glial subtypes, including astrocytes and ensheathing glia, infiltrate into the neuropil, forming tight associations with neuronal processes and synapses prior to eclosion at ∼96h APF^73,75^.

**Figure 1.**
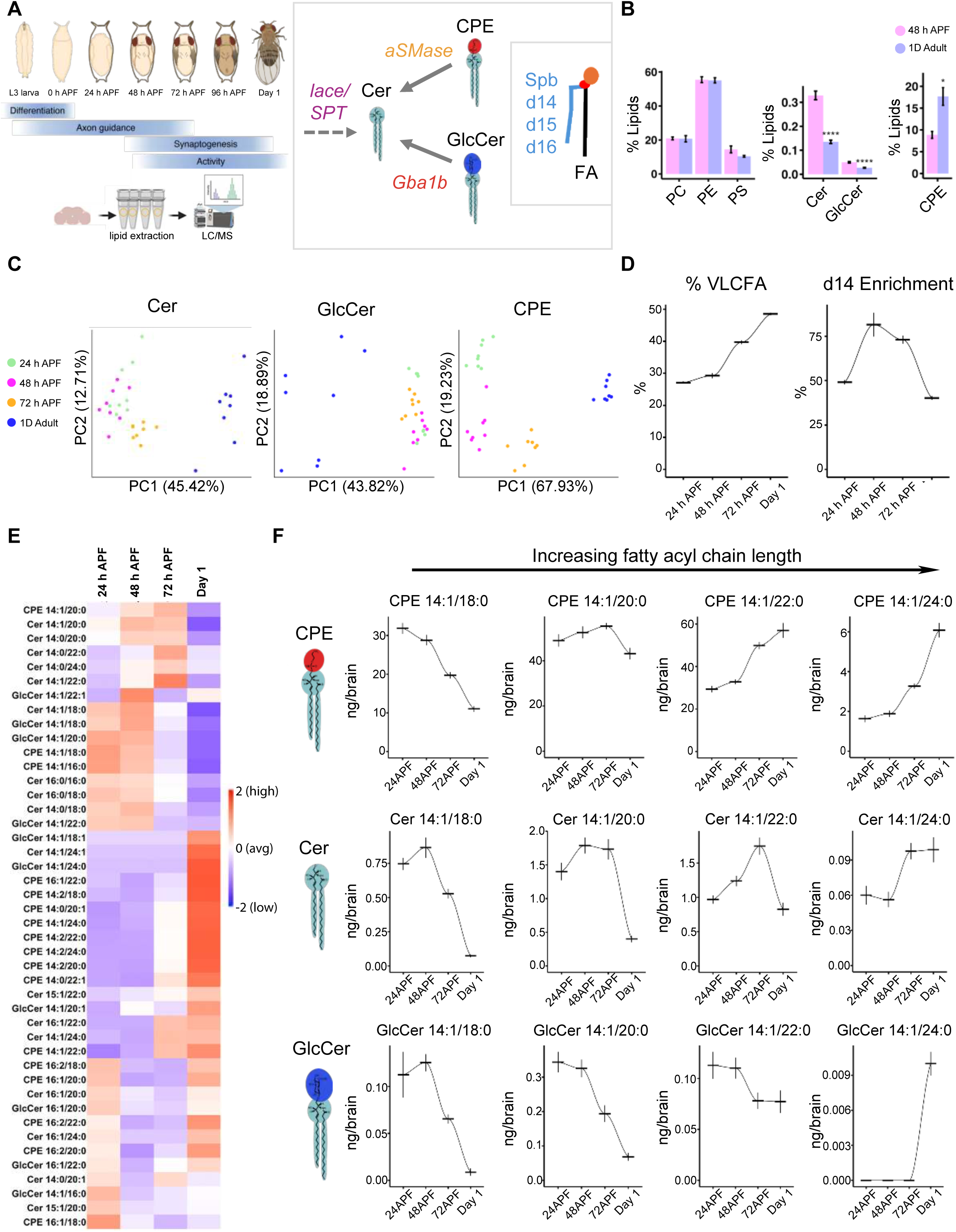
Sphingolipids are extensively remodeled during brain development. (A) Schematic of adult brain development in the fruit fly *Drosophila melanogaster*, and the major sphingolipid families Ceramide (Cer), Glucosylceramide (GlcCer), and Ceramide Phosphoethanolamine (CPE). (B) Mol% fraction of developing brain lipids (pink) taken at 48 hours after puparium formation (48h APF) compared to day 1 adult brains (blue). (C) Principal component analysis of the three sphingolipid classes across four developmental timepoints from dissected brains: 24h APF (green), 48h APF (magenta), 72h APF (orange), and newly eclosed adults (blue). (D) Very long-chain fatty acids (VLCFA) increase in late brain development, while d14 sphingoid base enrichment is transiently elevated in mid-development (E) Heatmap of sphingolipid species z-scored across developmental time (red = enriched). (F) Examples of major CPE, Cer, and GlcCer species that undergo developmental remodeling from shorter to longer acyl chains, quantified as ng/brain. Lipidomics in C-F represent 8 tubes of 15 brains per each timepoint, with mean and SEM plotted across time connected by loess lines in D. * p < 0.05, ** p < 0.01, *** p < 0.001, **** p < 0.0001

To quantify lipid levels during development, we first dissected control brains at mid-pupal development (48h APF), and across the first ten days of adult life, and used liquid chromatography-mass spectrometry (LC-MS/MS; see Methods) to measure the relative abundance of both membrane phospholipids and sphingolipids (Figure 1B, Figure S1). In line with previous studies of the adult brain^76–78^, three classes of membrane phospholipids, namely phosphatidylethanolamine (PE), phosphatidylcholine (PC), and phosphatidylserine (PS), represent a dominant fraction of the brain lipidome, and displayed little total change between mid-pupal and adult stages (Figure 1B). By contrast, all three major sphingolipid families, namely Ceramides (Cer), GlucosylCeramides (GlcCer), and Ceramide Phosphoethanolamine (CPE), displayed significant changes between these two stages, as Cer and GlcCer were ∼250-300% higher in pupal brains, while CPE levels were ∼50% lower in pupal brains (Figure 1B; Figure S1A-B). Thus, unlike the more common phospholipid classes, the total abundance of sphingolipids varied substantially during brain development.

Given these dramatic developmental changes, we next quantified specific sphingolipid species, including approximately 20 CPE species, 20 Cer species, and 10 GlcCer species that differed by chain lengths and saturations (Figure 1C). By convention, these lipid species are differentiated by head group (separating the three classes, Cer, GlcCer and CPE), and by the length and unsaturation of the two hydrophobic chains. For example, Cer 14:1/18:0 denotes a Ceramide containing a d14:1 sphingoid base comprised of one double bond and 14 carbons, paired with the fatty-acid stearic acid C18:0 (Figure 1A). Principal component analysis of sphingolipid distribution strongly separated 48h APF brains from all other adult ages (2,3,5, and 10 days), with developmental increases in 14:1/18:0 chains across all sphingolipids (Figure S1A-C). We did not observe major differences between sexes (Figure S1D).

Intrigued by this developmental sphingolipid signature, we measured lipids in brains from 24h APF, 48h APF, 72h APF, and newly eclosed adults (Figure 1C, Figure S1E). Principal component analysis of sphingolipid species strongly separated adult samples from pupal samples, accounting for ∼40-70% of variance across all three sphingolipid families (Figure 1C). While Cer and GlcCer species grouped all three pupal stages together, the CPE family cleanly separated all timepoints, suggesting that this lipid class undergoes particularly extensive developmental remodeling. By analyzing weighted averages of chain distributions across all three classes, we found that very long-chain fatty acids (VLCFA, defined as a fatty acyl tail longer than 20 carbons) increased during the second half of pupal development, and that d14 sphingolipids were transiently enriched at 48h APF and 72h APF (Figure 1D).

We next examined individual lipid species. The relative distribution of sphingolipids dynamically varied across time, with the most dramatic changes occurring between 72h APF and adulthood (Figure 1E; Figure S1F). Within each class of sphingolipids we observed a variety of patterns, with specific species that are relatively high during early pupal development and then fall, species that are low and then rise, and with some species that peaked at other pupal stages (Figure 1E-1F). Notably, across all sphingolipids, species with shorter fatty acyl chains were transiently abundant earlier in development, whereas longer chain species became enriched during late pupal development (Figure 1F). For example, 14:1/18:0 CPE continuously declined across development, whereas CPE 14:1/22:0 doubled between mid-pupal development and adulthood, and accounted for 35% of our total measured sphingolipid species at this timepoint.

Thus, brain development is characterized by highly dynamic sphingolipid remodeling, particularly during the latter half of pupal development, after neurons have chosen specific synaptic targets.

### Coordinated sphingolipid biosynthesis and catabolism constrain the developmental lipidome

How does the brain coordinately remodel sphingolipids? To gain insights into the genetic programs underlying sphingolipid remodeling during development, we examined the expression of major sphingolipid-regulating enzymes in longitudinal single-cell RNA sequencing datasets of the fly optic lobe^74^. We reasoned that sphingolipid remodeling could occur through either *de novo* biosynthesis via the serine palmitoyltransferase (SPT) complex located in the endoplasmic reticulum, or catabolic salvage via Glucocerebrosidase (GBA) or via acid Sphingomyelinase (aSMase) located in the lysosome. Intriguingly, both neurons and glia coordinately induce the expression of *lace* (SPTLCB2), the rate-limiting enzyme in the SPT complex, with neural expression peaking at 60h APF and glial expression at 48h APF (Figure 2A). In contrast, expression of *gba1b* (GBA) and *aSMase* (SMPD1) were restricted to glia, with *gba1b* transcripts peaking at 48-60h APF, the height of developmental GlcCer abundance, and *aSMase* peaking twice at 36h APF and 96h APF (Figure 2A). We validated the neuronal and glial expression of these three genes using *CRIMIC-GAL4* lines^79^, and found that *lace* was expressed in both neurons and glia, while *gba1b* and *aSMase* were exclusively expressed in glia (Figure S2A-S2B). These data suggest that compartmentalized glial catabolism and *de novo* biosynthesis in both neurons and glia could tune the developing brain sphingolipidome.

**Figure 2.**
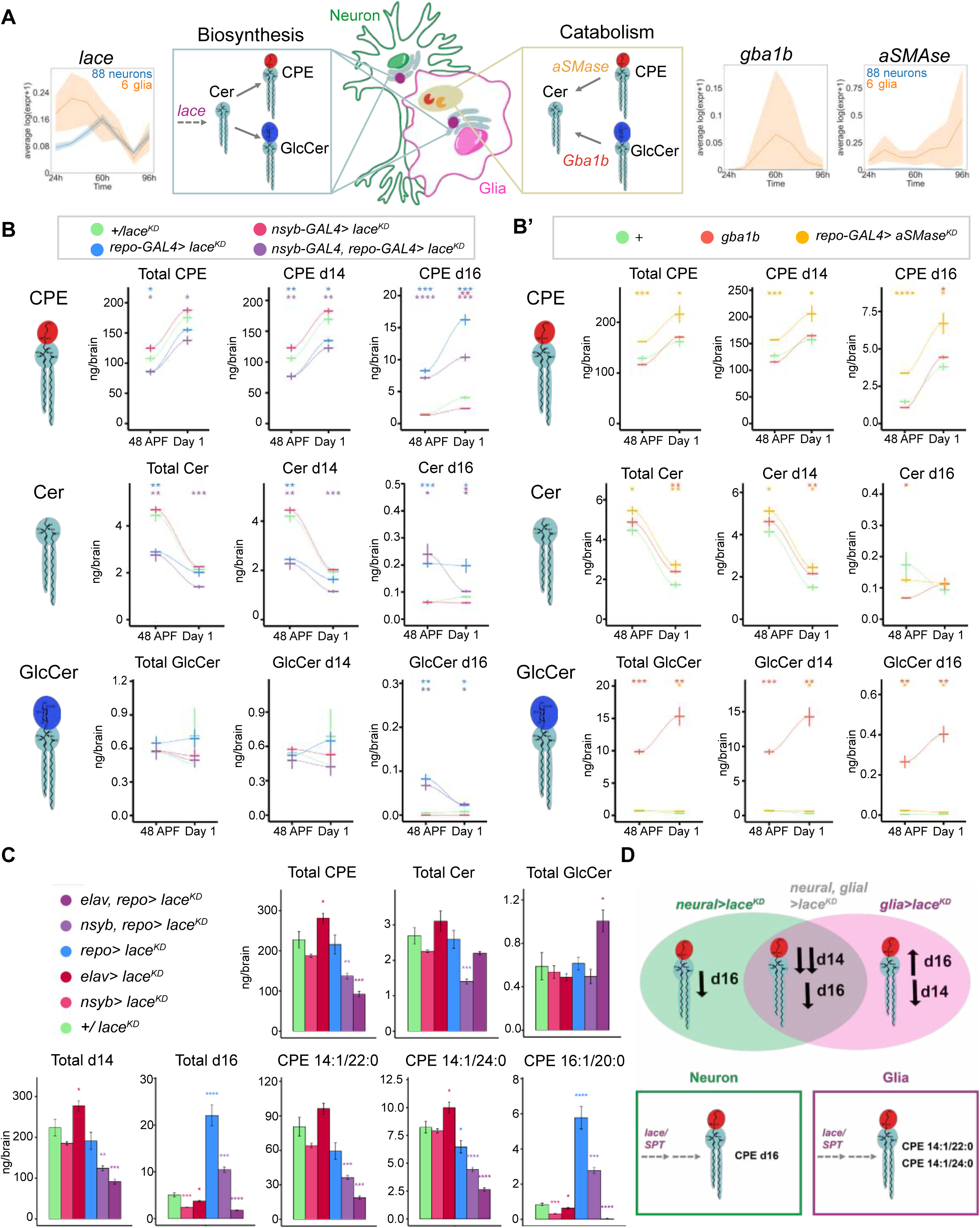
Coordinated sphingolipid biosynthesis and catabolism constrain the developmental lipidome. (A) Schematic of compartmentalized *de novo* biosynthesis via lace/SPT, and lysosomal catabolism mediated by Gba1b (cleaves GlcCer) and aSMase (cleaves CPE). RNA-sequencing data of these biosynthetic and catabolic genes are replotted from^74^, with the average expression of 88 neural clusters shown in blue, and six glial clusters shown in orange, from 24h to 96h APF. Data are represented as mean ± SEM. (B) Lipidomic analysis of total, d14, and d16 sphingolipids from controls (green) and brains depleted of *lace* via *RNAi* in glia (blue), neurons (red), or both (magenta) across two developmental timepoints, 48h APF and day 1. (B’) Lipidomic analysis of total, d14, and d16 sphingolipids from controls (green) and brains mutant for *gba1b^Δ^*(magenta) and *glial aSMase^KD^* (orange). (C) Lipidomics of control brains (green) versus *lace*-*RNAi (lace^KD^)* using early (*elav-GAL4*, dark red) and late neural (*nSyb-GAL4*, light red), glial (*repo-GAL4*, blue), or combined drivers (*elav-GAL4, repo-GAL4,* dark purple; and *nSyb-GAL4, repo-GAL4,* light purple). (D) Model for how removal of *lace* from neurons, glia, or both changes the brain sphingolipidome based on sphingoid base, with glial d14 and neuronal d16 contributions. n= 4 tubes of 15 brains per each timepoint, with mean and SEM plotted across time connected by loess lines or as barplots in D. * p < 0.05, ** p < 0.01, *** p < 0.001, **** p < 0.0001

To test this model, we first analyzed brain lipidomes from 48h APF and Day 1 adults in which *lace* was disrupted (Figure 2B; Figure S2C-S2D). We knocked down *lace* using RNAi in all glia with *repo-GAL4*, in all neurons with *nSyb*-*GAL4*, and in both classes with *nsyb-GAL4, repo-GAL4.* Consistent with neurons and glia both making biosynthetic contributions to sphingolipid levels, knockdown of *lace* in both cell types reduced the total levels of CPE and Cer across development, while total GlcCer levels were unaffected (Figure 2B). At this level, perturbations of *lace* in only neurons had modest effects, whereas knocking down *lace* in glia had larger effects on Cer and CPE. We next separated each sphingolipid class by the length of its sphingoid base, which can be d14 (typically representing approximately 90% of the total), d16 (representing 10% of the total), or trace levels of d15^80^. For all three sphingolipid classes, neuron-specific knockdown of *lace* had no effect on any d14 species, but reduced d16 CPE and Cer at Day1. Conversely, glial-specific knockdown of *lace* caused reductions in the levels of d14 CPE and Cer. Strikingly, however, the reductions in d14 CPE and Cer species seen in glial *lace* knockdown were paralleled by dramatically increased levels of d16 CPE and d16 Cer.

This reciprocity between neuronal and glial manipulations of *de novo* biosynthesis suggested that loss of biosynthesis in glia may trigger compensatory upregulation of biosynthesis in neurons. However, while glial *lace^KD^*brains displayed a three-fold increase in d16 CPE, simultaneously removing *lace* from neurons only incompletely reduced this effect (Figure 2B). Thus, d16 species may be produced in neurons before the *nSyb-*GAL4 mediated *RNAi* becomes effective, or they could be derived from non-neuronal (and non-glial) tissues. To discriminate between these two possibilities, we repeated these experiments using a second neuronal driver, *elav-GAL4*, that is expressed earlier in development. Strikingly, using *elav-GAL4, repo-GAL4* to drive *lace^KD^* in both neurons and glia completely suppressed the increased abundance of d16 species (Figure 2C). Moreover, d14 species were dramatically depleted, with CPE 14:1/22:0 and CPE 14:1/24:0 reduced to 25% of control levels (Figure 2C). Thus, while total sphingolipids are not reduced in glial *lace^KD^* due to compensatory increases in d16 balancing the loss of d14, removing *lace* from both neurons and glia prevented this compensation and reduced total sphingolipid levels to one third of controls (Figure 2D). This loss of sphingolipids had severe consequences for adult animals, who were largely unable to move. Thus, combined glial and neuronal *de novo* biosynthesis is required for an essential developmental sphingolipid bolus, with neuronal *lace* activity producing d16 sphingolipids, and both neuronal and glial *lace* producing d14 sphingolipids (Figure 2D).

We next used *gba1b* mutants to block GlcCer catabolism, and used glial knockdown of *aSMase* to block CPE catabolism, as *aSMase* null homozygotes are lethal^81^ (Figure 2B’; Figure S2C’-D’). Consistent with the expected biochemical selectivity, *gba1b* mutants specifically accumulate GlcCer, whereas in glial *aSMase^KD^* animals, CPE levels were substantially increased (Figure 2B’; Figure S). Both *gba1b* mutants and glial *aSMase^KD^* also had more modest effects on Cer levels, consistent with indirect effects on pathway flux. Taken together with the effects of the biosynthetic mutants, the developmental sphingolipidome is tuned in amplitude by both *de novo* biosynthesis from neurons and glia, and catabolism in glia.

### Disrupting sphingolipid levels alters endolysosomal processing and autophagy

We next sought to characterize the cellular consequences of perturbing sphingolipid catabolism and biosynthesis on brain development. Newly eclosed *gba1b* adults harbor hypertrophic lysosomes and, upon aging, accumulate p62/SQSTM1^63,65,82^, an adaptor protein that bridges polyubiquitinated substrates and autophagosomes^83^. We therefore predicted that catabolic mutants would dysregulate endolysosomal processing and accumulate p62. Consistent with this view, both *gba1b* and glial *aSMase^KD^* brains had enlarged lysosomes relative to control brains at 48h APF (Figure 3A-3B; Figure S3A-S3B). Similarly, the late endosome marker Rab7 accumulated in catabolic mutants within discrete neuropil boundary regions (Figure 3C). In addition, disrupting expression of *lace* in either neurons, glia, or both had no visible effects on either lysosomal enlargement or Rab7 accumulation (Figure 3B-C). However, contrary to our expectations, while catabolic mutants did not accumulate p62 at 48h APF, removing *lace* from neurons and glia caused large p62+ puncta accumulation at neuropil boundaries (Figure 3D). Thus, loss of sphingolipid catabolism during development causes endolysosomal defects, yet impairing sphingolipid biosynthesis in both neurons and glia causes autophagy defects.

**Figure 3.**
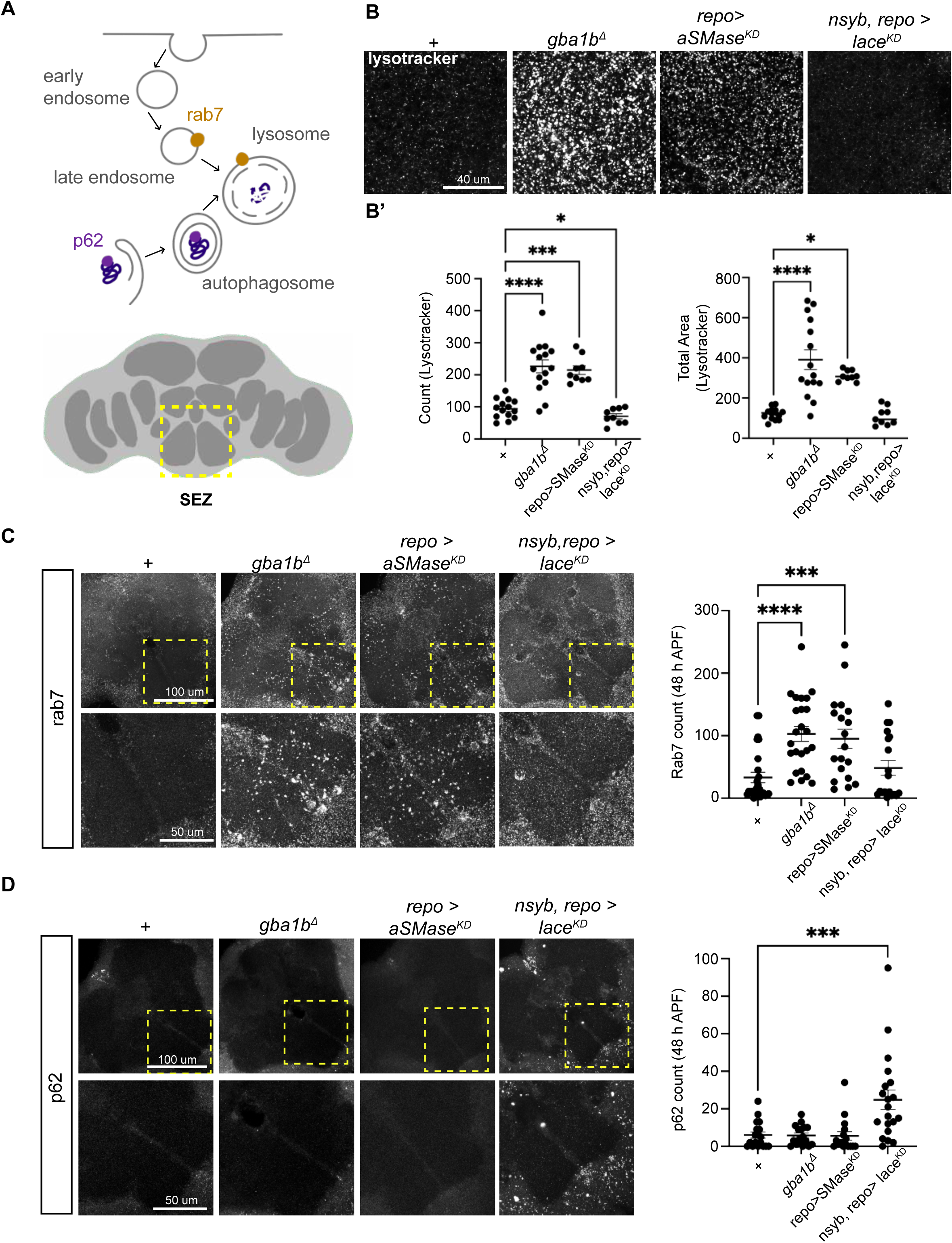
Disrupting sphingolipid levels alters endolysosomal processing and autophagy. (A) Schematic of endolysosomal and p62 autophagy pathways^26^, and the area of the brain selected for confocal imaging, the sub-esophageal zone (yellow box, SEZ). (B) Lysotracker labeling in max-intensity projections (MIP) of the central brain at 48h APF across control, catabolic, and biosynthetic mutants. Scale bar = 40 μm. (C) MIPs of Rab7 staining at 48h APF taken from control, catabolic, and biosynthetic mutants (upper panels, scale bar = 100 μm), with high magnification images of the SEZ (lower panels, scale bar = 50 μm). (D) MIPs of p62 staining at 48h APF, taken from control, catabolic, and biosynthetic mutants (upper panels, scale bar = 100 μm), with high magnification images of the SEZ (lower panels, scale bar = 50 μm). p62 aggregates appeared only following the dual depletion of *lace* in both neurons and glia using *nSyb-GAL4, repo-GAL4*. Data are represented as mean ± SEM. n > 10 brains per condition. * p < 0.05, ** p < 0.01, *** p < 0.001, **** p < 0.0001

### Neuronal and glial sphingolipid biosynthesis regulates glial autophagy

We next sought to determine where the autophagy defects caused by *lace* knockdown occurred in the brain. As observed during pupal development, newly eclosed adults in which *lace* was knocked down in both neurons and glia accumulated p62 (Figure 4A). The accumulation of p62 puncta near the edges of the neuropil was reminiscent of the positions of the cell bodies of astrocyte-like and ensheathing glia^84,85^. Using an antibody against Glutamine synthetase 2 (Gs2), which labels neuropil glia (including astrocyte-like and ensheathing glia (Figure S4A-S4C)^85^, we found that p62 accumulated specifically within Gs2+ glia, both inside the soma and within glial branches that extended into the neuropil (Figure 4A). Consistent with a broad defect in autophagy, these p62+ puncta also contained high levels of ubiquitin (Figure 4B). This phenotype was selectively observed when *lace* was removed from both neurons and glia, but not from either class individually, or in catabolic manipulations (Figure 4C). Thus, neuronal and glial sphingolipid biosynthesis is required for glial autophagy during development (Figure 4D).

**Figure 4.**
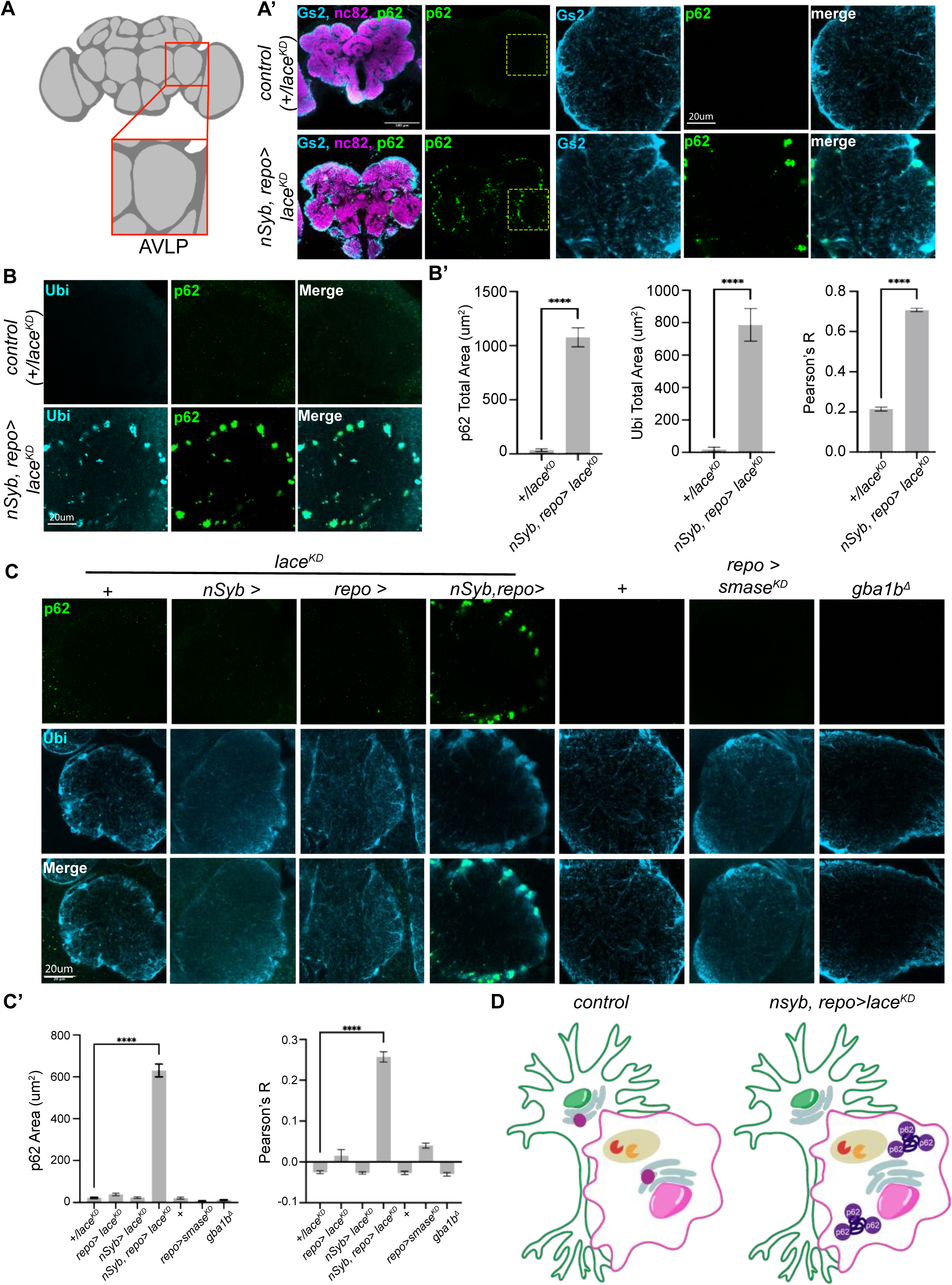
Neuronal and glial sphingolipid biosynthesis regulates glial autophagy. (A) Schematic of the adult fly brain and the anterior ventrolateral protocerebrum (AVLP) region selected for aggregate quantification. (A’) Confocal images of day 1 brains from controls and *lace-RNAi* (*lace^KD^*) in neurons and glia, stained for neuropil (nc82, magenta), p62 (green), and glutamine synthase 2 (Gs2, light blue). Scale bars = 100 μm; AVLP zoom scale bar = 20 μm. (B) Confocal images of the AVLP from day1 controls and *lace-RNAi* (*lace^KD^*) in neurons and glia, stained for neuropil (nc82, magenta), p62 (green), and ubiquitin (light blue). Scale bar = 20 μm. (B’) Quantification of p62 area, ubiquitin area, and the correlation between ubiquitin and p62. (C) AVLP stained with p62 (green) and Gs2 (light blue) across biosynthetic and catabolic mutants. p62 accumulates selectively in the simultaneous depletion of *lace* from neurons and glia using *nSyb-GAL4, repo-GAL4*. Scale bar = 20 μm. (C’) Quantification of p62 and p62-Gs2 colocalization from AVLP in C. (D) Model for p62 aggregation in glia upon simultaneous loss of *lace* (purple circle in ER) from neurons and glia. Data are represented as mean ± SEM. n > 10 brains per condition. * p < 0.05, ** p < 0.01, *** p < 0.001, **** p < 0.0001

### CPE is cell-autonomously required for autophagy in glia

To identify the sphingolipids required to sustain glial autophagy, we depleted various sphingolipid biosynthetic enzymes in neurons, glia, or both (Figure 5A-B; Figure S5). Depletion of either the Ceramide Synthase, *schlank (*CerS), or the CPE synthase *cpes* in all glia caused p62 accumulation within Gs2+ glia (Figure 5B). Conversely, neuronal depletion of *schlank* or *cpes* did not (Figure 5B). Moreover, knocking down *cpes* expression only in Gs2+ glia using two independent drivers was sufficient to trigger p62 puncta (Figure 5C; Figure S5A), pointing to a cell-autonomous role for CPE synthesis. Strikingly, while *cpes* trans-heterozygous null animals also harbored p62 aggregates, this phenotype was strongly rescued by *Gs2-GAL4* driving expression of UAS-Cpes constructs, whereas neuronal drivers failed to rescue, as did most other glial drivers (Figure S5C-E). Thus, Gs2+ glia autonomously require CPE biosynthesis for autophagy.

**Figure 5.**
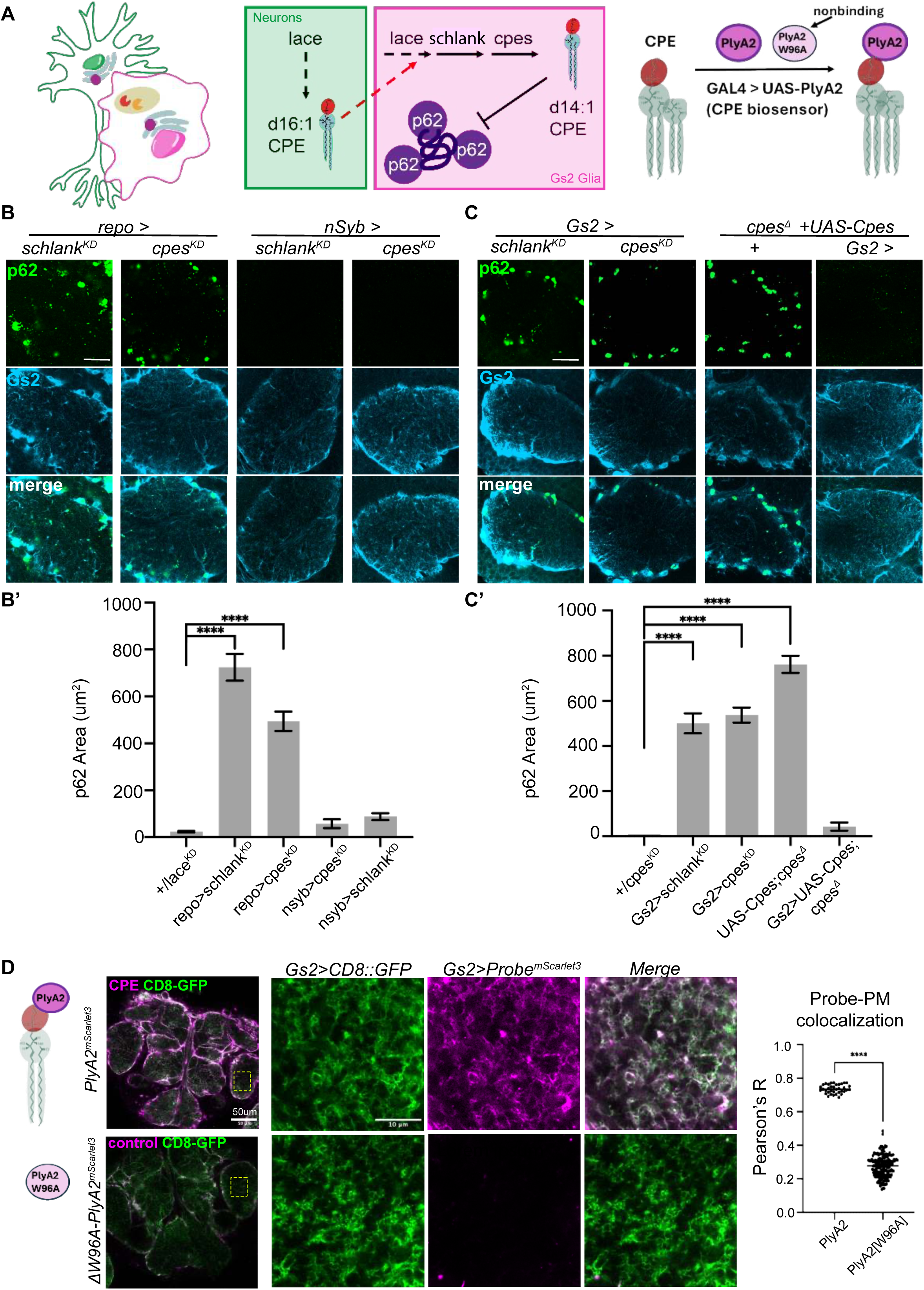
CPE is cell-autonomously required for autophagy in glia. (A) Model depicting the requirement for *lace* expression in neurons and glia, or *schlank* (CerS) and *cpes* (CPE synthase) expression in glia. To label CPE lipids, PlyA2 domain conjugated at the C-terminus to mScarlet3 is expressed using *GAL4/UAS*. PlyA2 selectively binds CPE, but the W96A mutation abrogates binding while still expressing mScarlet3 as a control. (B) AVLP stained with p62 (green) and Gs2 (light blue) in neuronal (*nSyb-GAL4*) or glial (*repo-GAL4*) knockdown of *schlank* or the CPE synthase *cpes*. Scale bar = 20 μm. B’, quantification of p62 area in AVLP. (C) AVLP stained with p62 (green) and Gs2 (light blue) in Gs2+ glial knockdown of *schlank* or *cpes* using *Gs2-GAL4.* The *cpes* null p62 phenotype is rescued by *Gs2-GAL4* driving *UAS-Cpes*. Scale bar = 20 μm. C’, quantification of p62 area in AVLP. (D) Genetically encoded *PlyA2::mScarlet3* biosensor for CPE expressed in Gs2+ glia (magenta), with CD8-GFP (green) labeling glial membranes. The control probe expresses *PlyA2[W96A]:mScarlet3*, a mutated version of PlyA2 that abrogates lipid binding. Zooms into AVLP show fine membrane processes labeled by GFP that also accumulate the CPE biosensor but not the control probe. Scale bar = 50 μm and 10um for zooms into AVLP membranes. Data are represented as mean ± SEM. A-C, n > 10 brains per condition. D, n = 5 brains. * p < 0.05, ** p < 0.01, *** p < 0.001, **** p < 0.0001

We next examined whether the headgroup of CPE was required for its function in glia, using heterologous expression of human *Sphingomyelin Synthase* enzymes hSMS1 or hSMS2. While flies do not normally produce sphingomyelin endogenously given the absence of genomic SM synthases^86^, introducing hSMS1 into *cpes* nulls generates ectopic Sphingomyelin, while heterologously expressing hSMS2 generates both CPE and ectopic Sphingomyelin^87^. Intriguingly, expressing either hSMS2 or hSMS1 in Gs2+ glia in *cpes* null mutant animals strongly rescued the p62 phenotype, suggesting that Sphingomyelin can functionally replace CPE in neuropil glia for autophagy (Figure S5B, Figure S5E).

Finally, to test if Gs2+ glia contain CPE, we generated a genetically encoded CPE probe using the non-cytolytic pleurotolysin domain PlyA2 fused to mScarlet3 at the C-terminus. Previously, using purified liposomes and *ex vivo* assays, PlyA2 was demonstrated to selectively bind to the sphingolipids CPE or SM, but not to identical headgroups of the glycerophospholipids phosphatidylethanolamine or phosphatidylcholine^88–92^. Thus, PlyA2 should label CPE in *Drosophila*, which lack SM and SM synthases^86^. We also generated a negative control for this probe, *PlyA2^W96A^*, which abolished CPE binding in liposomes and in CNS explants^88,89^. To detect CPE in Gs2+ glia, we co-expressed *CD8::GFP* with either *PlyA2::mScarlet3* or PlyA2^W96A^*::mScarlet3*. Under these conditions, glia in newly eclosed animals robustly accumulated PlyA2 labeling, but did not accumulate PlyA2^W96A^ (Figure 5D). Thus, Gs2+ glia contain CPE at membranes in close contact with neuronal synapses.

Taken together, these results demonstrate that CPE synthesis in Gs2 glia is necessary and sufficient to prevent the accumulation of p62 aggregates. Crucially, as knockdown of *lace* in neurons and glia phenocopied the loss of CPE production specifically in glia (achieved via knockdown of *cpes* and *schlank*), glia can use sphingolipid precursors produced in both neurons and glia to generate CPE (Figure 5A).

### Transfer of neuronal lipids to glia by the phagolysosome

How can neuronally derived sphingolipids be used to make CPE in glia? We hypothesized that glia might obtain lipids via phagocytosis of neuronal membranes. Consistent with this hypothesis, we observed that the MEGF10 homolog *draper*^93^ was enriched in p62 positive puncta when *lace* was removed from both neurons and glia, or when *cpes* was removed from either Gs2+ glia or in *cpes* null animals (Figure 6A). This phenotype was rescued in *cpes* null animals when *Cpes* was expressed only in Gs2+ glia (Figure 6A). To test this idea further, we labeled glial membranes with *mCD8::GFP* and neuronal membranes with *myr::TdTomato* and then removed *cpes* from Gs2+ glia. Under these conditions, we observed glial membranes enclosing neural membranes, subsets of which were decorated with ubiquitin (Figure S6A).

**Figure 6.**
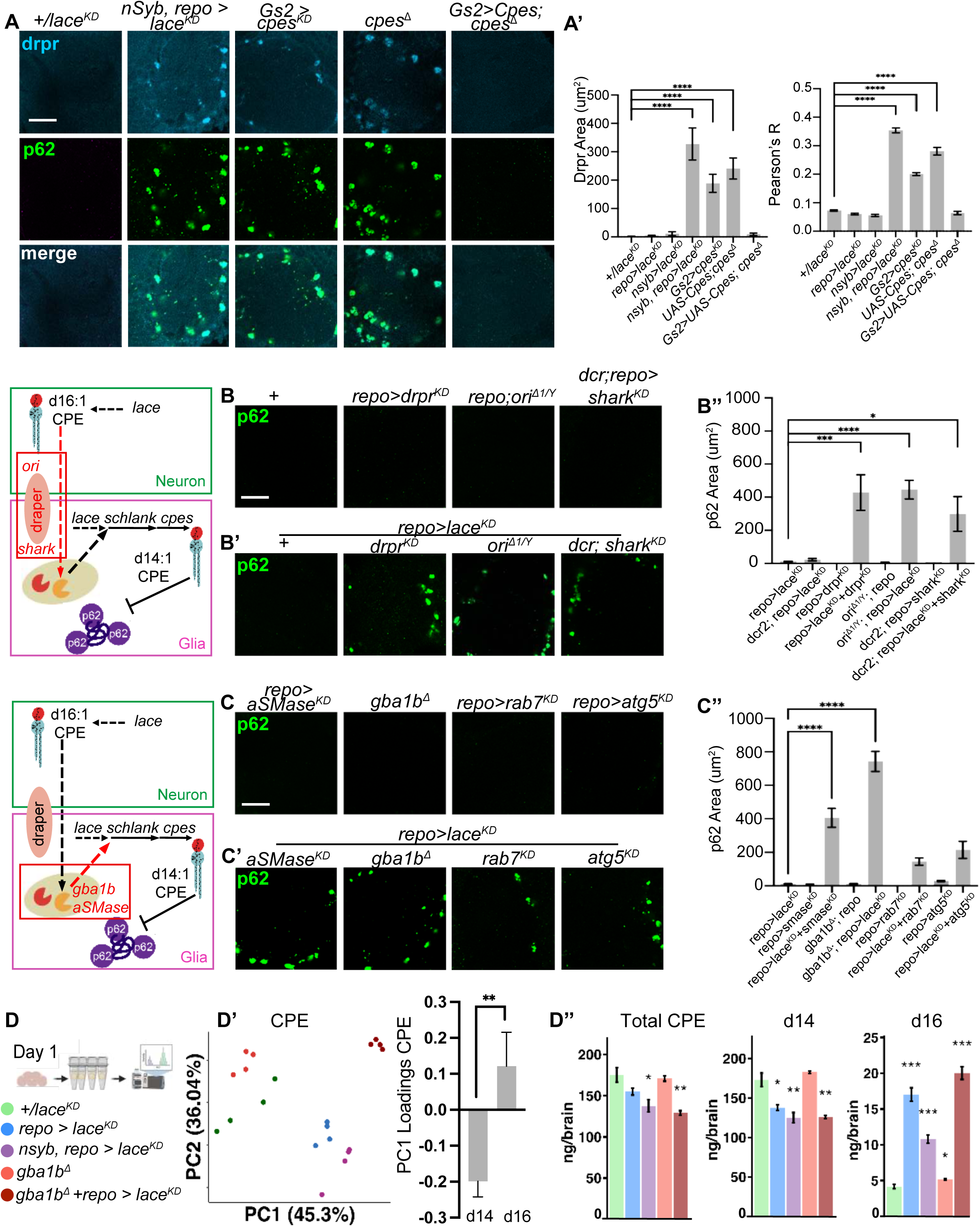
Transfer of neuronal lipids to glia by the phagolysosome. (A) The phagocytosis receptor draper (drpr, light blue) accumulates in p62+ aggregates (green) in CPE-depleted Gs2 glia (using *nsyb-GAL4, repo-GAL4* to deplete *lace*, or *Gs2-GAL4* to deplete *cpes*). *cpes* nulls also accumulate drpr but are rescued by *Gs2-GAL4* expressing *UAS-Cpes*. Scale bar = 20 μm. (A’) Quantification of drpr accumulation and colocalization with p62 from B. (B) AVLP stained with p62 (green) in glial knockdown of phagocytosis genes *drpr* or *shark* using *repo-GAL4,* and in males mutant for *ori*. Scale bar = 20 μm. (B’) AVLP stained with p62 (green) from brains depleted of *lace* in glia using *repo-GAL4* combined with the simultaneous removal of phagocytosis genes *drpr* or *shark,* and in males mutant for *ori*. Scale bar = 20 μm. (B’’) Quantification of p62 accumulation from D-D’. (C) AVLP stained with p62 (green) in lysosomal catabolic perturbations (*gba1b^Δ^* or glial knockdown of *aSMase* using *repo-GAL4*), or autophagy/late endosome protein knockdown of *atg5/rab7* in glia using *repo-GAL4*. Scale bar = 20 μm. (C’) AVLP stained with p62 (green) from brains depleted of *lace* in glia using *repo-GAL4* combined with the simultaneous removal of lysosomal genes (*gba1b^Δ^* or glial knockdown of *aSMase* using *repo-GAL4*), or autophagy/late endosome protein knockdown of *atg5/rab7* in glia using *repo-GAL4*. Scale bar = 20 μm. (D’’) Quantification of p62 accumulation from E-E’. (D) Lipidomics of day 1 brains from controls (*+/lace^KD^*, green), glial *lace* knockdown (*repo-GAL4*, blue), neural+glial *lace* knockdown (*nSyb-GAL4, repo-GAL4,* light purple), *gba1b^Δ^* mutants (red), and *gba1b^Δ^* mutants in glial *lace* knockdowns (*gba1b^Δ^* +*repo-GAL4>lace^KD^,* dark red). (D’) PCA plot for CPE of genotypes in F, with PC1 loadings summed for d14 vs d16. (D’’) Total CPE, total d16, and total d14 sphingolipids from genotypes in F, in ng/brain. Data are represented as mean ± SEM. n > 10 brains per condition for A-C, n = 4 tubes of 15 brains for D. * p < 0.05, ** p < 0.01, *** p < 0.001, **** p < 0.0001

If glia obtain sphingolipids either through lace-dependent *de novo* biosynthesis or by phagocytosing and recycling neuronal sphingolipids, then simultaneous blockade of both mechanisms should impair CPE production and lead to p62 accumulation. We therefore combined knockdown of *lace* in glia with phagolysosome perturbations. Indeed, combining knockdown of *lace* in glia with knockdown of the phagocytosis receptor *draper,* its bridging chemokine *orion*^94^, or the downstream effector kinase *shark*^95^ triggered p62 accumulation (Figure 6B-B’’; Figure S6B-S6E). Similarly, targeting lysosomal recycling by removing *gba1b* or *aSMase* together with glial *lace* knockdown also caused accumulation of p62 (Figure 6C-C’’). Importantly, removal of these catabolic enzymes or phagocytosis effectors alone did not lead to p62 accumulation (nor did knockdown of *lace* in glia alone). This surprising genetic interaction between biosynthesis (*lace*) and catabolism (*draper, gba1b, aSMase*) points to a key role for glial salvage of neuronal lipids for future biosynthesis. Indeed, lipidomics of *gba1b* mutants combined with glial knockdown of *lace* (*gba1b^Δ^* +*repo>lace^KD^*) revealed strong effects on CPE lipids, with PC1 strongly separating this genetic interaction from controls or plain *gba1b* mutants (Figure 6D-D’). Similar to the simultaneous loss of *lace* from both neurons and glia, the separation of *gba1b^Δ^*+*repo>lace^KD^*from controls was driven by loss of d14 and gain of d16 CPEs. Indeed, total CPE and d14 sphingolipids were depleted in *gba1b^Δ^*+*repo>lace^KD^*, yet the d16 sphingolipids that become elevated in glial knockouts of *lace* remained highly enriched (Figure 6D’’). These data are consistent with Gba1b and lysosomal catabolism functioning downstream of the phagocytosis of neuronal d16 sphingolipids to produce d14 CPE for glia.

### CPE is required for glial infiltration and synapse numbers

We next investigated the developmental consequences for Gs2+ glia deprived of CPE. During the late stages of brain development, astrocyte processes grow into synaptic neuropils by first extending large primary branches, and then by infiltrating fine secondary processes towards synapses^73,96^. We therefore tested if either primary branch formation or subsequent secondary infiltration requires CPE. To do this, we sparsely labeled Gs2+ glia using SPARC^97^ in controls and in animals where *cpes* had been knocked down using *Gs2-GAL4*. We quantified astrocyte morphology across the late stages of pupal development (Figure 7A; Figure S7A). At 48h APF, astrocytes in control brains and *cpes^KD^* brains were indistinguishable, and displayed little outgrowth (Figure 7B). At 72h APF, control astrocytes grew substantially by extending primary branches and increasing both branch number and surface area. At the same stage, astrocytes lacking *cpes* also grew, extending primary branches that had reduced complexity. At eclosion, when control astrocytes displayed elaborate branching and substantial growth in surface area, *cpes* deficient astrocytes were morphologically aberrant, with strongly reduced branching complexity and reduced surface area (Figure 7B; Figure S7B). To visualize the outgrowth of fine processes more closely, we examined cross sections of the neuropil and quantified the number of discrete glial segments (Figure 7C). Strikingly, loss of *cpes* increased the thickness of the primary branches relative to controls while simultaneously reducing the degree of discrete glial infiltrations (Figure 7C’). Thus, during brain maturation, synapse-infiltrating Gs2+ astrocytes undergo dramatic membrane growth and branching, morphological changes that require CPE.

**Fig.7.**
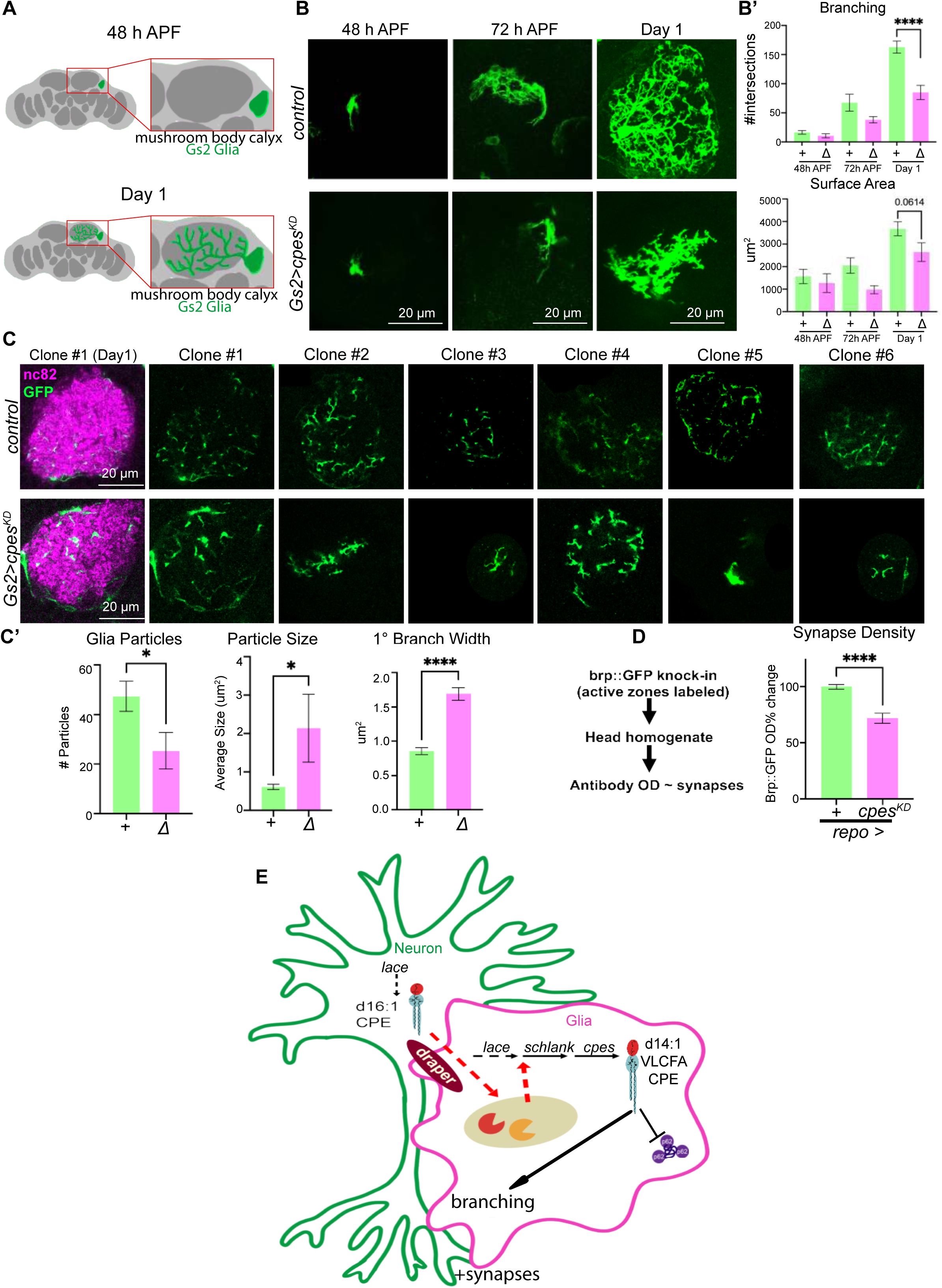
CPE is required for glial infiltration and synapse numbers. (A) Schematic of the mushroom body calyx region used for quantification of glial morphology as glia infiltrate synapses between 48h APF and eclosion (day 1). (B) Sparsely labeled Gs2+ astrocytes expressing CD8-GFP were imaged across brain development in controls and *Gs2-GAL4* mediated *cpes* knockdowns at 48h APF, 72h APF, and day 1. Shown are maximum intensity projections of clones in the mushroom body calyx. Scale bars = 20 μm. n = 5 clones for 48h APF, n = 8 clones for 72h APF, n = 20 clones for day 1. (B’) Quantification of branching (sholl analysis) and surface area across controls (green) and *cpes* knockdowns (magenta). (C) Cross-sections of mushroom body calyx clones from controls (top rows) or *cpes* knockdowns (bottom rows) at day 1, with CD8-GFP (green) marking glial membranes in the synaptic neuropil (magenta). Scale bars = 20 μm. n = 15 cross sections. (C’) Quantification of cross-section glial particle number and size, and primary branch width across controls (green) and *cpes* knockdowns (magenta) at day 1. n = 15 cross sections. (D) Synapse protein levels were measured using an ELISA assay to detect Brp::GFP^99^ in fly head lysates from glial *cpes* knockdowns with *repo-GAL4* (magenta) normalized to controls (green). n = 10 replicates (20 heads total). (E) Model. Data are represented as mean ± SEM. * p < 0.05, ** p < 0.01, *** p < 0.001, **** p < 0.0001

As astrocytes can regulate the development and maintenance of synapses^98^, we asked if this disruption of glial infiltration was associated with a loss of synapses. By using a quantitative ELISA-based assay to measure expression of the synaptic active zone protein Bruchpilot (Brp)^99^, we found that Brp levels were reduced when *cpes* was knocked down in glia (Figure 7D). Moreover, we observed qualitatively similar reductions in Brp staining in the mushroom body calyx (Figure S7C), where we also observed changes in Gs2 glial morphology when *cpes* was cell-autonomously knocked down.

Finally, we profiled lipids from controls and brains depleted of *cpes* in Gs2 glia, which account for ∼3.5% of the entire brain used for LC-MS. Despite the sparsity of this genetic manipulation, we detected reductions in VLCFA CPEs 14:1/22:0 and 14:1/24:0 (Figure S7D). Consistent with a requirement for VLCFA sphingolipids, Gs2 glia depleted of the Ceramide Synthase *schlank* were rescued from autophagy defects via heterologous expression of human CerS2 (Figure S7E-E’). In contrast, hCerS6, which selectively produces C14-C16 sphingolipids, could not rescue autophagy defects (Figure S7E-E’). However, while hCerS2 localized extensively throughout glial processes, hCerS6 localized more selectively to primary branches (Figure S7F), meaning that either shorter chain lengths or altered localization could underlie the failed rescue. Nonetheless, these data show that VLCFA sphingolipids are sufficient for autophagy in Gs2 glia, and that VLCFA CPE levels are reduced when *cpes* is selectively removed from this cell class.

Taken ogether, these results argue that VLCFA CPE is required for glia to infiltrate the neuropil as circuits mature, a process that is essential to adult synapse regulation (Figure 7E).

## Discussion

### Sphingolipids play a central role in circuit maturation in the developing brain

Assembling the complex architecture of the brain requires precisely coordinated changes in neuronal and glial morphologies, changes that necessitate spatiotemporally controlled reorganization of lipid membranes^100,101^. While many lipids play structural roles, lipids can also act as signals in the developing brain. Here we define a new role for lipids during development, whereby a precisely timed bolus of sphingolipids governs circuit maturation, enabling glia to infiltrate the developing brain after neurons have chosen synaptic partners. We show that a specific subclass of sphingolipids, long-chain CPE species, are essential for both autophagy and membrane elaboration in neuropil glia that must extend fine processes throughout the synaptic neuropil. Strikingly, glia bypass a blockade in *de novo* sphingolipid biosynthesis by consuming neuronal sphingolipids to sustain glial infiltration. Disruption of this metabolic coupling prevents glial infiltration into the developing brain, leading to synaptic defects. Thus, combining a timed developmental bolus of sphingolipid production with a non-autonomous lipid-based interaction between neurons and glia creates a coordinating mechanism by which neuronal and glial membranes can be remodeled concurrently, at the correct time.

### Coordinating glial infiltration, CPE production, and synaptic activity

Long-chain CPE sphingolipids accumulate in the brain from 72h APF into adulthood (Figure 1), corresponding to when Gs2+ astrocytes ramify processes (Figure 7B) that contain CPE at plasma membranes (Figure 5D). Crucially, loss of glial CPE reduces synapse density (Figure 7D) and caused photosensitive epilepsy, pointing to the importance of this glial lipid in the nonautonomous control of neuronal functions^59^. Intriguingly, electrical activity begins and increases throughout this developmental period^72,73^, and work in many species has described activity-dependent induction of glial phagocytic remodeling of synapses^75,102,103^. Consistent with this model, Draper/MEGF10 is elevated during late pupal development and mediates early-life circuit remodeling^75,103,104^. At the same time, hyperactivation of Draper or phagocytosis causes degeneration^62,105–107^, showcasing that membrane rearrangements must be carefully balanced. Taken together, neuronal activity may couple glial phagocytosis to enable CPE production and infiltration, an exciting possibility for future investigation.

### Sphingolipid biosynthesis regulates glial autophagy

One of the surprising results of our work is the finding that sphingolipids are critical regulators of autophagy in glia. Disrupting sphingolipid biosynthesis in both neurons or glia, or blocking CPE production autonomously in neuropil glia, results in the accumulation of p62 aggregates (Figures 3-5). Intriguingly, blocking sphingolipid catabolism also leads to the accumulation of p62 protein aggregates in older brains^65,82^, hinting that impaired endolysosomal processing may ultimately lead to defects in sphingolipid biosynthesis, downstream of impaired catabolism. Consistent with this hypothesis, blocking *gba1b* when glia are also deprived of sphingolipid biosynthesis caused a dramatic autophagy defect (Figure 6E), demonstrating that catabolism serves a crucial anabolic function through the liberation of sphingolipid precursors.

Why does removing CPE from neuropil glia cause defects in autophagy? CPE/SM trafficked from the plasma membrane via endolysosomal pathways could control endomembrane rearrangements relevant to the timely processing of autophagosomes or phagolysosomes. For example, SM selectively partitions into intraluminal vesicles from endosomal compartments when visualized by super-resolution microscopy^108^, and CPE is critical for multivesicular body formation at the cytokinetic furrow^89^. Thus, specific pools of CPE could directly control autophagy through sorting to intraluminal vesicles and generating Ceramides essential for autophagy-lysosomal flux^109–111^, such as during Schwann cell myelinophagy^112^. Regardless of the specific regulatory mechanism, studies of bulk lipidomics from flies also revealed strong ceramide induction during pupal development^113^, and timed increases in sphingolipid levels occur during brain development in mice^87,114^ and humans^115,116^. Thus, across many species, sphingolipids may tune autophagic flux both within and outside the nervous system.

### Developing glia play a central role in sphingolipid metabolism

Our findings and recent work on the dihydroceramide desaturase^117^ support a model where glia play a central role in sphingolipid biosynthesis. In this vein, primary chick oligodendrocytes have 4-fold higher SM synthase activity than primary neurons, and a correspondingly 4-fold higher SM/Ceramide ratio^118^. Similarly, iPSC-derived astrocytes and microglia have higher rates of *de novo* sphingolipid biosynthesis than motor neurons, and only glia responded transcriptionally when ceramide synthesis was blocked^119^. Interestingly, these transcriptional changes included prominent alterations in axon guidance pathways in astrocytes, perhaps reflecting the fact that glia can grow processes alongside pathfinding axons in the developing brain^120^. Whether these gene expression changes relate to our observed glial morphological defects would be interesting to explore in future work, and broadly support the notion that the substantial role of glia in regulating both sphingolipid biosynthesis and catabolism is evolutionarily ancient.

Our results with cell-type specific *lace* manipulations also point to robust compensatory networks between neurons and glia that can rescue the loss of biosynthetic capacity in one cell type. In particular, we observed reciprocal changes in d14 and d16 sphingoid bases following the depletion of *lace* in only glia. Intriguingly, clones of *lace* cells induced in developing epithelia were rescued from endocytic trafficking defects when surrounded by neighboring wild-type cells^121^, and large, but not small, *SPT* neural clones caused wiring phenotypes^56^, pointing to nonautonomous rescue by the transfer of lipids or enzymes. Moreover, astrocytes can compete for phospholipids when growing into the adult brain^122^, where bulk lipid transfer permits early developmental phagocytosis of neuronal debris^123^. Although extracellular vesicles and lipoproteins represent possible cellular mechanism for transporting lipids from one cell to another^70,124^, here we show that glial phagocytosis of neuronal membranes endows glia with a mechanism to bypass loss of *de novo* biosynthesis. Thus, when sphingolipids begin to decline in *lace* depleted glia, increased production (and/or consumption) of neuronal membranes can generate sufficient d14 CPE to sustain glial morphogenesis and autophagy.

What molecular mechanism underlies this compensatory interaction? One possibility is that the ORMDL family of SPT negative regulators that are acutely sensitive to Ceramide levels could sense and respond to the loss of sphingolipids in one cell-type^125–127^. As excess sphingolipid production in oligodendrocytes caused by ORMDL inactivation drives myelination defects, balanced sphingolipid production is critical for glial membrane lipid composition and function^128^. Similarly, neurons can compensate for the loss of Sphingosine-1-phosphate Lyase by increasing catabolism while decreasing biosynthesis^129^. Thus, a tightly controlled developmental bolus of sphingolipids is essential for brain development, and the brain can robustly correct cell-type deficits in lipid production by intracellular and intercellular means.

### Conserved functions for long-chain CPE/SM in glial membrane morphogenesis

Sphingolipids are induced during PNS myelination during the early postnatal period^114^, and CNS myelination encompasses a dynamic shift towards long-chain C24 sphingolipids in oligodendrocytes^87^. Similarly, cultured astrocytes, microglia, and oligodendrocytes contain more SM than neurons^34^, with weighted sphingolipid chain lengths varying by cell type^34^. Indeed, oligodendrocytes are replete with both sphingomyelin and Galactosylceramide^49^. In vertebrates, C22 and C24 long-chain sphingolipids are generated by Ceramide Synthase 2 coupled to specific elongases^130,131^, and *Cers2*-deficient mice have myelination phenotypes^52^. As *Drosophila* only harbor a single ceramide synthase^132^, the molecular mechanism that permits the shift to long-chain sphingolipids in the developing fly brain remains unclear. Although fly brains contain myelin-like wrappings^133^, they lack canonical myelin compaction proteins and the major myelin lipid GalCer. However, GalCer shares some similar biochemical properties with CPE, including high phase transition temperatures and the formation of tubules *in vitro* that are sensitive to chain length^134–136^. Intriguingly, CPE is required for peripheral glial wrapping^60^, adult cortex glia morphogenesis^59^, and neuropil glial ramification (this work). Moreover, the requirement for CPE in neuropil glia autophagy can be rescued by heterologous production of sphingomyelin using glial-specific expression of human sphingomyelin synthases (Fig. S5B). Thus, the highly ramified and thin glial membranes in both flies and vertebrates may require functionally comparable lipid species.

How might long-chain CPE/SM support dynamic membrane rearrangements to drive glial infiltration? SM accounts for ∼45% of exoplasmic lipids quantified in red blood cells, while inner leaflets only contained ∼2% SM^137^. This asymmetry likely exists for glial CPE, as the Cpes active site that catalyzes the CDP-alcohol phosphotransferase reaction faces the lumen of the Golgi^86^. Thus, SM and CPE may broadly support plasma membrane morphology and rearrangements at the exoplasmic leaflet, perhaps by sculpting lipid and protein microdomains^138,139^ and/or by regulating accessible cholesterol^140^. It is notable that 35% of the detected adult fly brain sphingolipidome was 14:1/22:0 CPE, which features an 8 carbon asymmetry that could enable lipid tail interdigitation and trans-bilayer coupling across the plasma membrane^141^. This long-chain sphingolipid may be particularly important for the specialized thin membranes required to fully infiltrate developing synapses.

Our observations of sphingolipids as regulators of glial infiltration into the brain have intriguing parallels in other cell types. For example, in *C. elegans*, the anchor cell invades a specific epithelial layer during vulval morphogenesis, an invasive process that is sphingomyelin-dependent; in fly larval sensory neurons, developmental ramification requires long-chain CPE^142^; and in the vertebrate immune system, macrophage engulfment of client particles also requires long-chain sphingomyelins^143^. Taken together with our findings that CPE production is required for glial infiltration into the synaptic neuropil, long-chain CPE/SM sphingolipids may play an evolutionarily ancient role in regulating plasma membrane rearrangements across many cell types.

## Acknowledgments

Research done at Stanford is conducted on Muwekma Ohlone land, and research done at UCSF is conducted on Ramaytush Ohlone land. We thank members of the Clandinin and Vaughen labs for discussions of the manuscript, and Estela Stevenson for excellent technical assistance provided to the Clandinin lab. We thank Tobi Stork and Jaeda Coutinho-Budd for discussion and sharing flies; Jairaj Acharya for discussing CPE; Richard Stanley for the p-Shark antibody; and Rushika Perera, Andrew Yang, and Leanne Jones for access to confocal microscopes. Funding includes the Sandler PBBR program (JPV), the Simons Foundation (TRC), NIH NINDS R21 NS12458 (TRC), NINDS K99 NS133298 (TRJ), and the Stanford Interdisciplinary Graduate Fellowship (EKT). GK is supported by the intramural division of the NCI, NIH, HHS. We gratefully acknowledge the Bloomington Drosophila Stock Center (NIH P40OD018537), the Harvard TRiP Center (R24 OD030002)^144^, and the Stanford Vision Core (P30EY026877).

## Author Contributions

Conceptualization: EKT, TRC, JPV

Methodology: EKT, IMRS, RJL, TTN, GK, TRJ, MTC, TRC, JPV

Investigation: EKT, IMRS, RJL, GK, TRJ, JPV

Visualization: EKT, RJL, TRC, JPV Funding acquisition: VM, MTC, TRC, JPV

Project administration: VM, MTC, TRC, JPV Supervision: MTC, TRC, JPV

Writing – original draft: EKT, TRC, JPV Writing – review & editing: EKT, TRC, JPV

## Resource Availability

### Lead contact

Further information and requests for resources and reagents should be directed to and will be fulfilled by the lead contact, John Vaughen

### Materials availability

Flies and plasmids are available from the lead contact, and flies will be deposited at the Bloomington Drosophila Stock Center (BDSC).

### Data and code availability

Sphingolipid lipidomics are deposited as Supplemental Table 3. All FIJI and R scripts are available upon request. All data and information required to analyze the data in this paper are available from the lead contact upon request.

### Inventory of Supplemental Information

***Figures S1-S7 (7 figures)***

***Table S1, detailed genotypes***

***Table S2, sequences of PlyA2 and PlyA2-W96A plasmids***

***Table S3, lipidomics***

## Materials and Methods

### STAR Methods

#### KEY RESOURCES TABLE

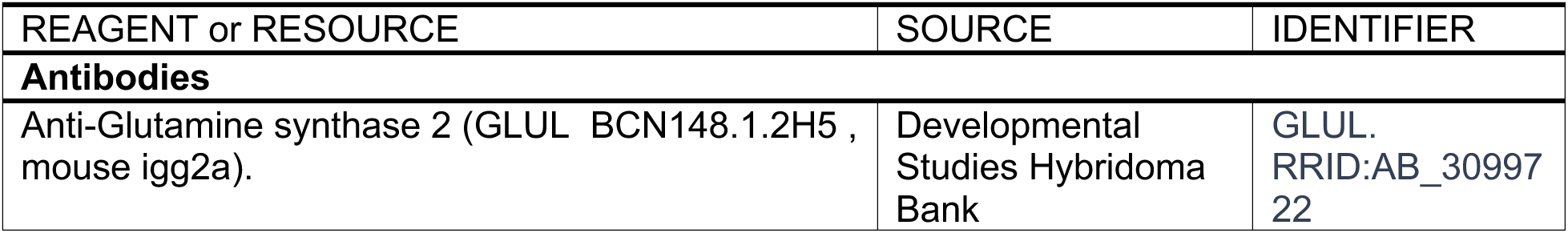
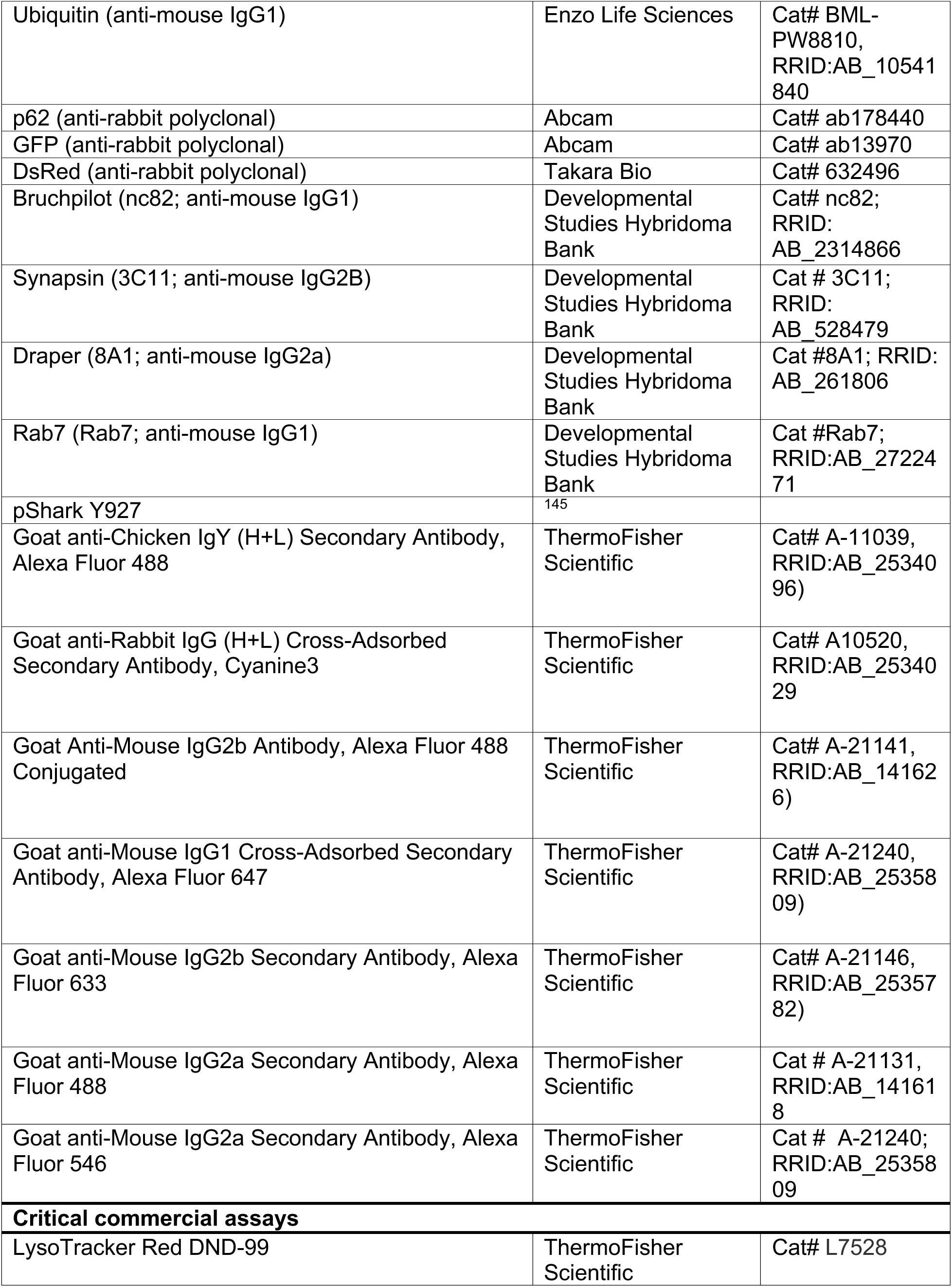
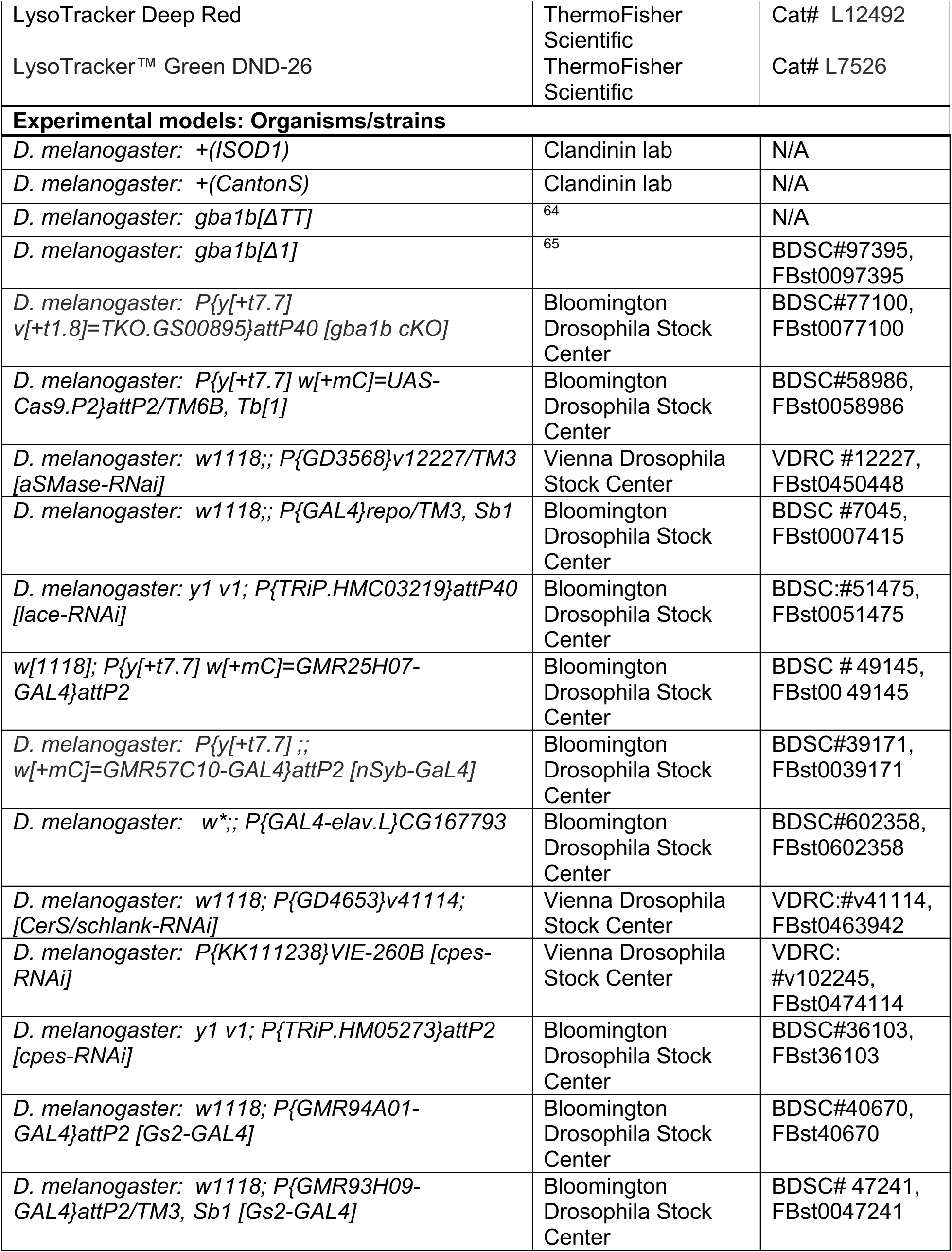
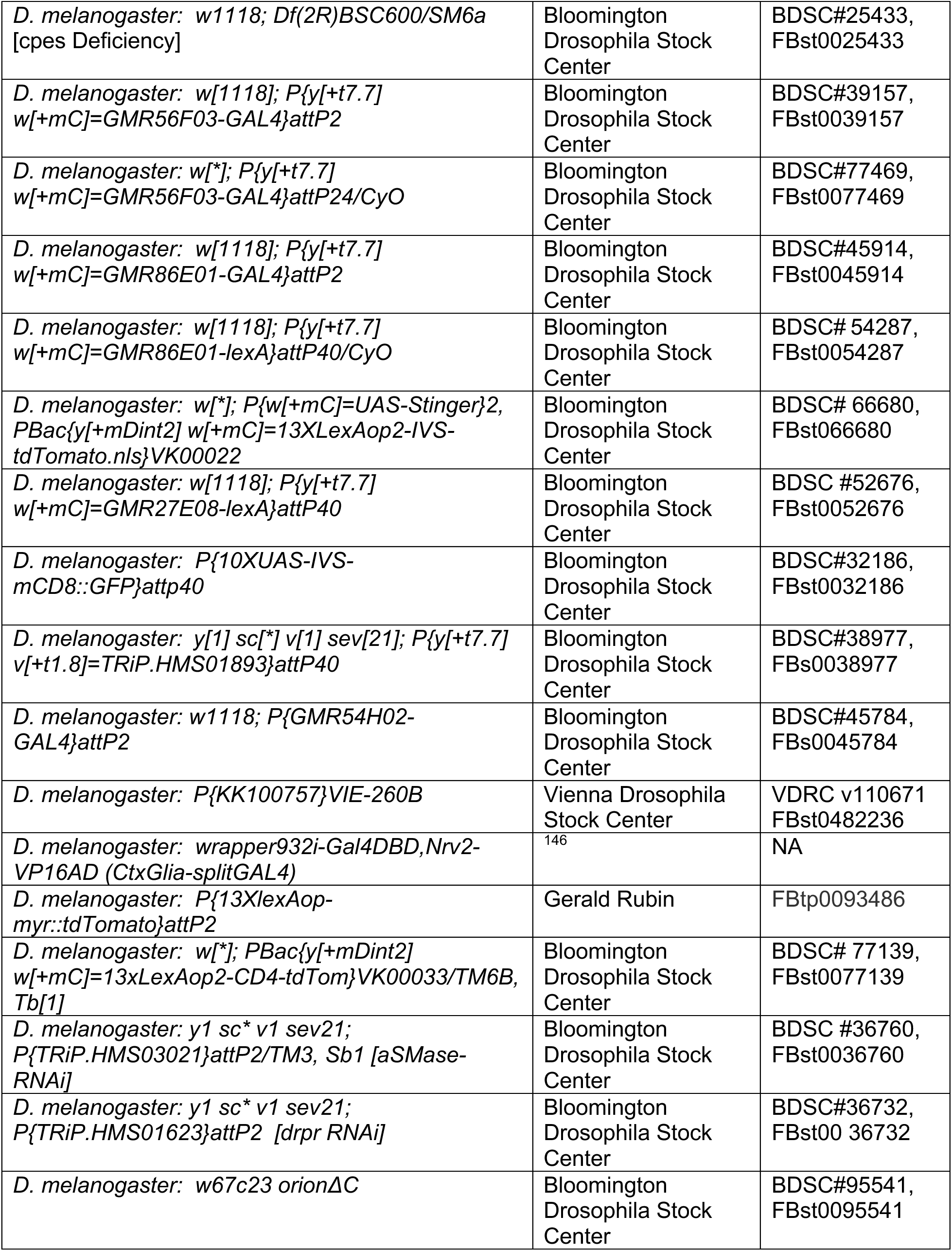
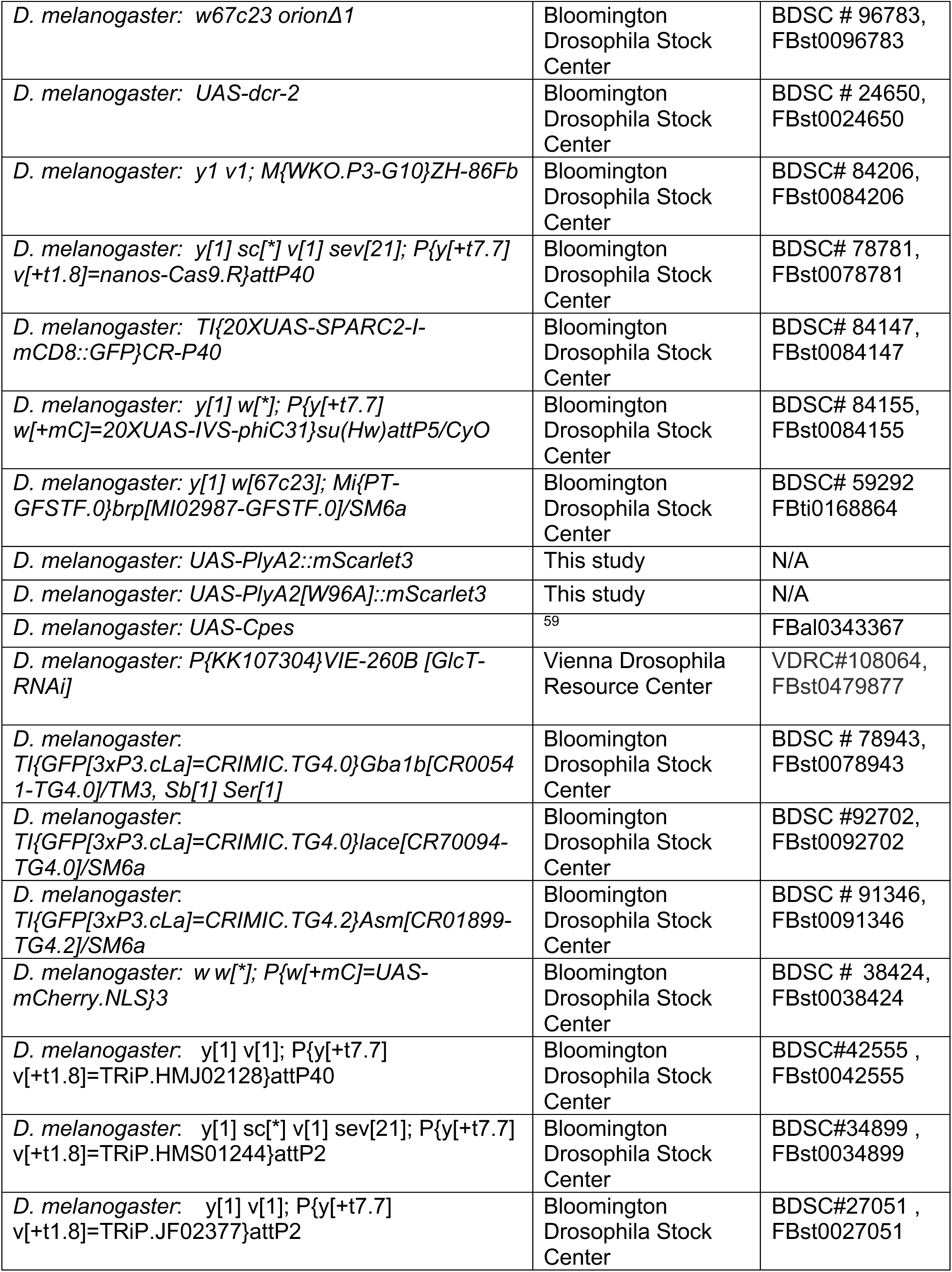
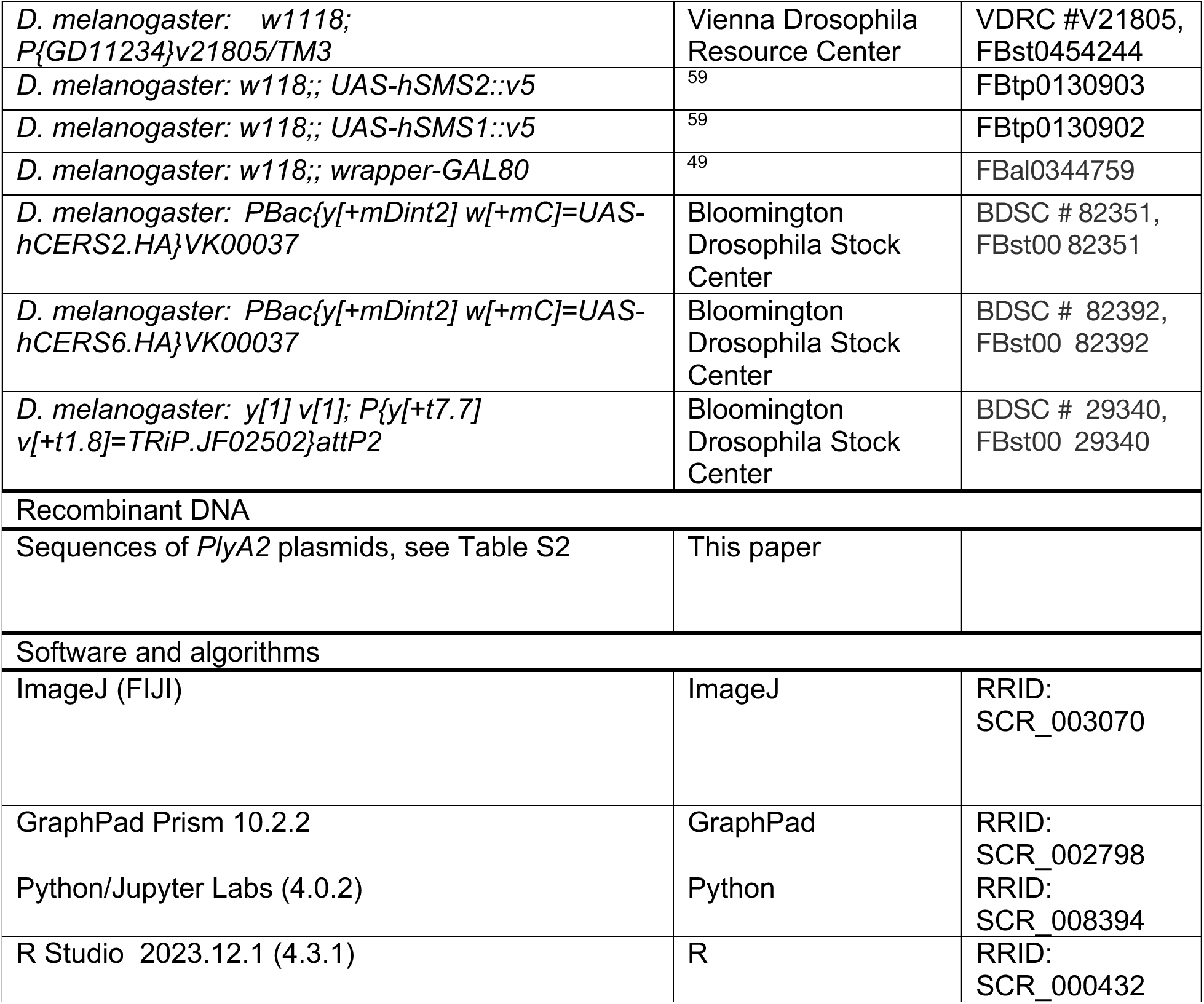

##### *Drosophila* Maintenance

Flies were maintained on standard molasses and cornmeal food (R food, LabExpress). Experiments were maintained in light/dark (LD) incubators for 12:12 light:dark cycles at 25°C with 50%-70% relative humidity. *RNAi* experiments for the genetic interaction with *repo>lace-RNAi* were conducted at 29°C, all other experiments were conducted at 25°C.

###### Molecular Genetics

We use FlyBase to find information on stocks, phenotypes, and sequences^147,148^. The PlyA2 sequence MAYAQWVIIIIHNVGSKDVKIVNLKPSWGKLHADGDKDTEVSASKYEGTVIKPDEKLQINACGRS DAAEGTTGTFDLVDPADGDKQVRHFYWDCP**W**GSKANTWTVSGSNTKWMIEYSGQNLDSGAL GTITVDTLKKGN was directly fused to mScarlet3 at the c-terminus, and cloned into pJFRC7-20XUAS vector (addgene #26220) using codon optimization for *Drosophila melanogaster* by Twist Bioscience. Plasmids were injected by Bestgene Inc and integrated into *attp40* (2^nd^ chromosome) or *attp2* (3^rd^ chromosome) by phiC31 integrase mediated recombination.

###### Brain Explant Imaging (LysoTracker and PlyA2)

Flies in batches of 5 were dissected under cold 1X dissection saline (103mM NaCl, 3mM KCL, 5mM TES, 1mM NaH_2_PO_4_, 4mM MgCl) and placed in Terasaki plates containing 1X dissection saline before individual transfer to 12µL of freshly diluted LysoTracker solution (1:500 dilution from stock LysoTracker solutions, 2µM, Thermofisher) for 2 minutes before immediate transfer to 100µL of saline on a microscope slide. Brains were pressed to the bottom of the saline bubble and oriented with dorsal side up (apposed to the coverslip).

Brains were imaged on a Leica SP8 confocal using a 40X Lens (N.A. 1.30) at 3X digital zoom. Z-stacks of 5 slices through the cortical region were acquired from optic lobes. Batches of 5 brains (interleaving control and experimental brains) were transferred to individual wells of Saline, LysoTracker, and individual slides in parallel. For imaging *myrTdT* and *GFP* membranes and *PlyA2* sensors, a similar live preparation (omitting Lysotracker) was followed. PlyA2 was imaged on a Zeiss LSM900 with Plan-APOCHROMAT 40X (1.3 NA) Oil lens with 0-6x digital zoom, or a LSM980 63X lens with 0-4x digital zoom with airyscan.

###### IHC Dissections and Staining

As with LysoTracker, flies were anesthetized on ice and immobilized in dissection collars. The proboscis was removed under cold dissection saline and then freshly diluted 4% paraformaldehyde (PFA) was added to the dissection collar (32% EM grade PFA (EMS)). Brains were fixed for 25 minutes; after 5 minutes of immersion in 4% PFA, the remaining head cuticle and surrounding fat was gently removed. Post-fixation, brains were washed three times in 1X PBS before completing the dissection in collars and removing brains into Terasaki wells with 0.5%PBSTx (TritonX-100 0.5% in 1X PBS). Brains were permeabilized for 30 mins then transferred to blocking solution (10%NGS in 0.5%PBSTx) for 40 minutes before adding primary antibodies in 0.5%PBSTx+10%NGS (1:10 CSP, 1:200 anti-FK2 polyubiquitin, 1:500 anti-p62 (Abcam ab178440), 1:10 bruchpilot (nc82, DSHB), 1:10 Glutamine Synthase 2 (GLUL, DSHB), 1:10 Rab7 (DSHB), 1:10 synapsin (3C11, DSHB), pShark (1:100), and 1:10 draper (8A1, DSHB) for 24-48 hours at 4°C. Brains were then washed three times in 0.5%PBSTx before transfer to appropriate secondaries (1:500, Thermo Fisher Scientific) for 2-4 hours. Brains were washed three times in 0.5%PBSTx before placing in 70% glycerol for clearing, then mounted and imaged in Vectashield. Images were collected using a Leica SP8 confocal microscope equipped with a 40X lens (N.A. 1.30) at 3X digital zoom. For AVLP p62 staining, images were collected on a Zeiss LSM900 with Plan-APOCHROMAT 40x/1.3 NA Oil lens with 2X digital zoom, with Z-stacks of 15 slices through the AVLP acquired to quantify aggregates.

The following antibodies were obtained from the Developmental Studies Hybridoma Bank, created by the NICHD of the NIH and maintained at The University of Iowa, Department of Biology, Iowa City, IA brp (nc82) developed by E. Bruchner^149^; draper 8A1 developed by M. Logan^150^; Rab7 developed by S. Munro^151^; SYNORF (Synapsin) developed by E. Bruchner^152^; and GLUL (developed by CDI labs)^153^.

###### Image analysis

Confocal files were imported into FIJI and analyzed with custom macros for particles for ubiquitin, lysotracker, or p62, or macros for colocalization, as previously reported^65^. For analysis of Gs2+ astrocyte branching, max-intensity projections of SPARC clones from the mushroom body calyx (collected at 3x digital zoom on 40x lens with 0.29 μm z-step resolution) were analyzed by sholl analysis (from ROI defined around the soma; start radius 4 μm, step size 2 μm, end radius 40 μm). The FIJI 3D Manager was used to calculate surface area from these clones. Particle analysis scripts were modified to analyze glial segments from cross-sections of the mushroom body calyx. Metrics were exported to excel and analyzed by ANOVA corrected via Tukey’s test for multiple comparisons in Prism GraphPad.

###### Lipidomics

15 brains per condition (genotype/timepoint) were dissected in quadruplicate (4 separate tubes). Dissections were in 1X dissection saline (see “IHC dissections” above). Newly eclosed flies were anesthetized on ice; all non-brain tissue (fat/hemocytes, larger trachea) was removed, as well as the retina (entirely). Single dissected brains were immediately transferred to Eppendorf tubes containing 20µl saline on ice; after 15 brains were added (10-15mins), 180µl of methanol was added (90% methanol v/v) and brains were snap-frozen on dry ice and stored at −80°C until further analysis. Brains were analyzed for sphingolipids and phospholipids as previously reported using a LC triple quadrupole MS^65^. Ng/Brain and % relative class brain data were analyzed, FDR corrected for multiple comparisons, and plotted in R studio.

###### ELISA

Flies of the indicated genotypes were collected at 3 days post-eclosion, frozen on dry ice and stored at −80°C. Heads were dissociated from flies by vortexing and 2 heads were combined in each sample. Samples were homogenized and an ELISA assay performed against GFP as previously described^89^. Optical density values for each sample were normalized to the control sample average. Male and female flies were normalized separately within each sex due to baseline differences in Brp levels in females and males. Samples were run across 3 independent experiments.

## Supplemental Figures and Figure Legends

**Figure S1, related to Figure 1.**
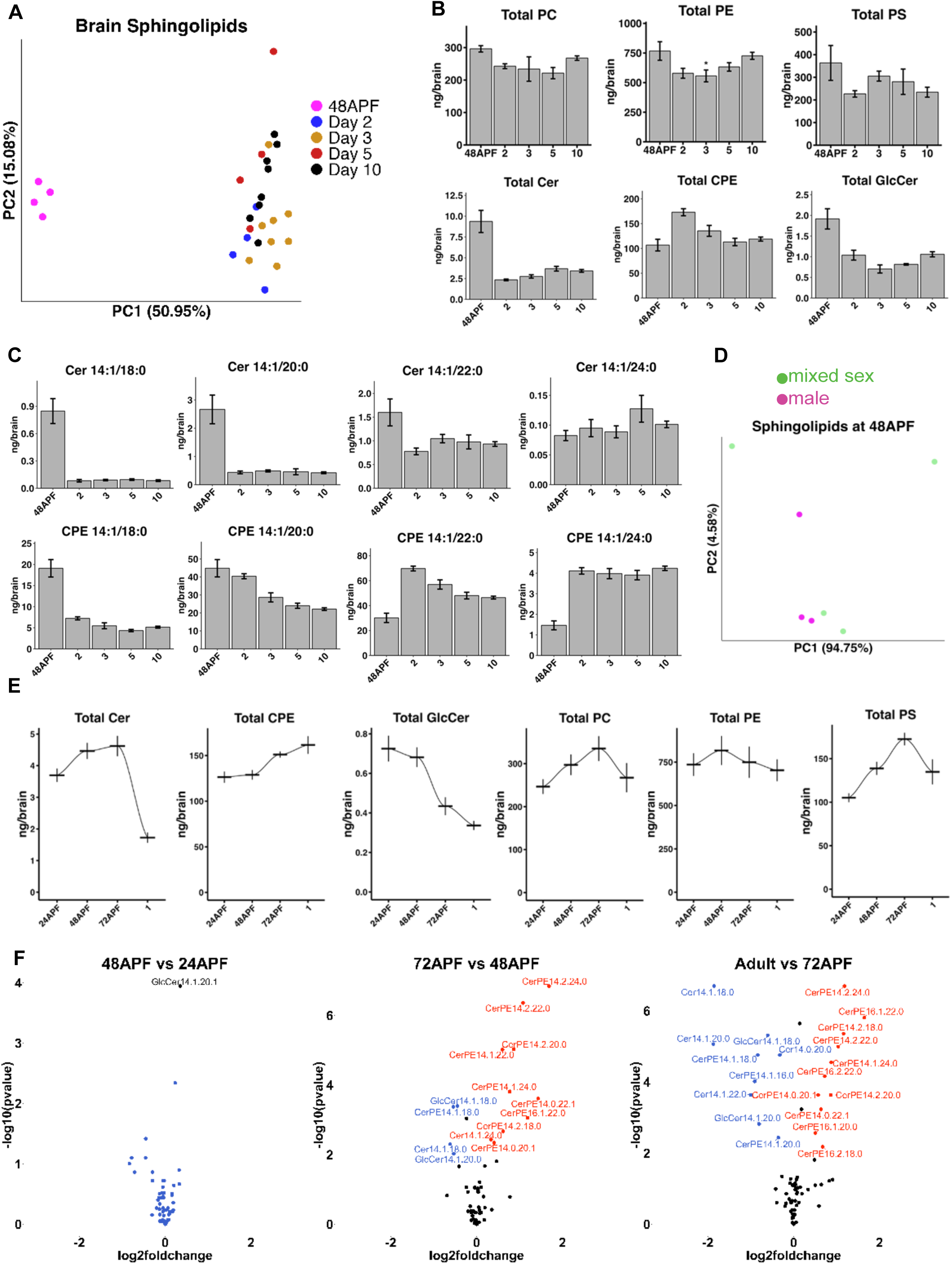
(A) PCA analysis of sphingolipids from control brains dissected during development and 48h APF (magenta) and adult ages. (B) Total phospholipids and sphingolipids measured at 48h APF and multiple adult ages, quantified in ng/brain. (C) Developmentally modulated sphingolipid levels in ng/brain at 48h APF and adult ages. (D) PCA analysis at 48h APF of male (magenta) and mixed sex (green) brains. (E) Developing brain total sphingolipids and phospholipids (ng/brain). (F) Volcano plots between the 4 developmental timepoints (48h APF versus 24h APF; 72h APF versus 48h APF; and day 1 versus 72h APF). Lipidomics in E-F represent 8 tubes of 15 brains per each timepoint, lipidomics in A-D are 8 tubes of 15 brains per timepoint for day 3 and day 10, and 4 tubes of 15 brains per timepoint for all other ages. Data are represented as mean ± SEM. * p < 0.05, ** p < 0.01, *** p < 0.001, **** p < 0.0001

**Figure S2, related to Figure 2.**
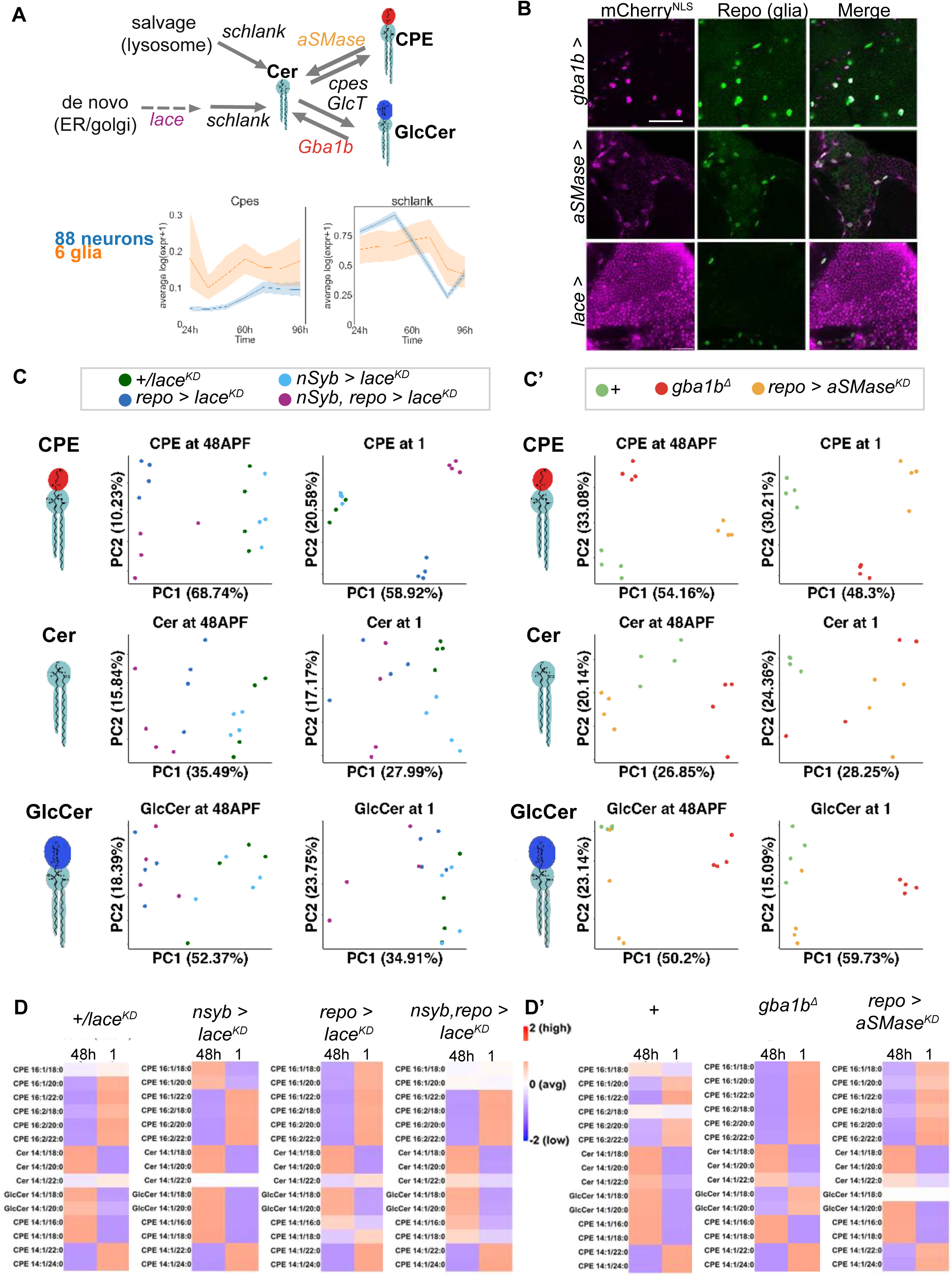
(A) Simplified diagram of enzymes in sphingolipid biosynthetic and salvage (catabolic) pathways. scRNAseq data replotted from^74^, 88 neural cluster average is shown in blue, and glial cluster average depicted in orange, from 24h to 96h APF. Data are mean ± SEM. (B) *CRIMIC-GAL4* driven *nls-mCherry* using *Gba1b-GAL4*, *aSMase-GAL4,* or *lace-GAL4* (magenta), co-stained with the glial marker repo (green). *Gba1b-*expressing cells were 97% repo+, *aSMase-*expressing cells were 91% repo+, and *lace-*expressing cells were 8% repo+ (n = 5 brains each genotype). Scale bar = 20µm. (C) PCA analysis of CPE, Cer, and GlcCer at 48h APF and day 1 for biosynthetic manipulations targeting *lace* by glial (*repo-GAL4*, blue), neural (*nSyb-GAL4*, light red), or combined neural and glial drivers (*nSyb-GAL4, repo-GAL4,* light purple) versus controls (*+/lace-RNAi*, light green). (C’). PCA analysis of CPE, Cer, and GlcCer at 48h APF and day 1 for catabolic manipulations (*gba1b^Δ^*, red) and *aSMase* knockdown in glia (*repo-GAL4 >* SMase^KD^, orange), versus controls (green). (D-D’) Z-scores of major developmentally regulated sphingolipids across time in biosynthetic (D) and catabolic (D’) manipulations reveals that the global patterns of developmental changes in sphingolipids is relatively robust across genotypes. n= 8 tubes of 15 brains per each timepoint for C-D. * p < 0.05, ** p < 0.01, *** p < 0.001, **** p < 0.0001

**Figure S3, related to Figure 3.**
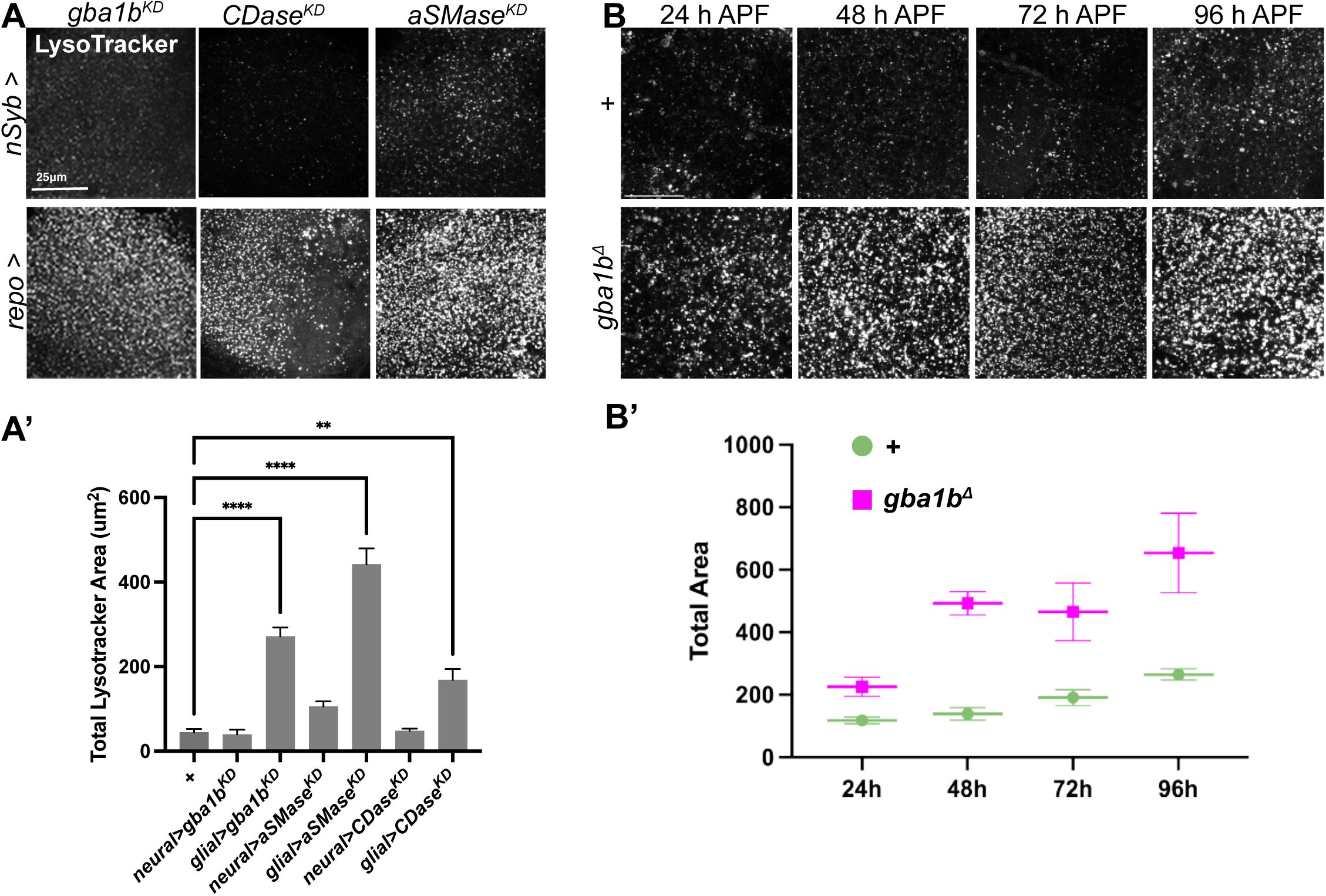
(A-A’) Confocal images of lysotracker staining (white), with maximum intensity projections of optic lobes from neural (*nSyb-GAL4*) or glial (*repo-GAL4*) knockdowns of sphingolipid catabolic enzymes. A’, quantification of lysotracker area. Scale bar = 25µm. (B’) Timecourse of lysotracker from central brains in *gba1b^Δ^* and control brains across pupal development, quantified in B’ (green = control, magenta = *gba1b^Δ^*). Scale bar = 20µm. Data are represented as mean ± SEM. n > 10 brains all experiments. * p < 0.05, ** p < 0.01, *** p < 0.001, **** p < 0.0001

**Figure S4, related to Figure 4.**
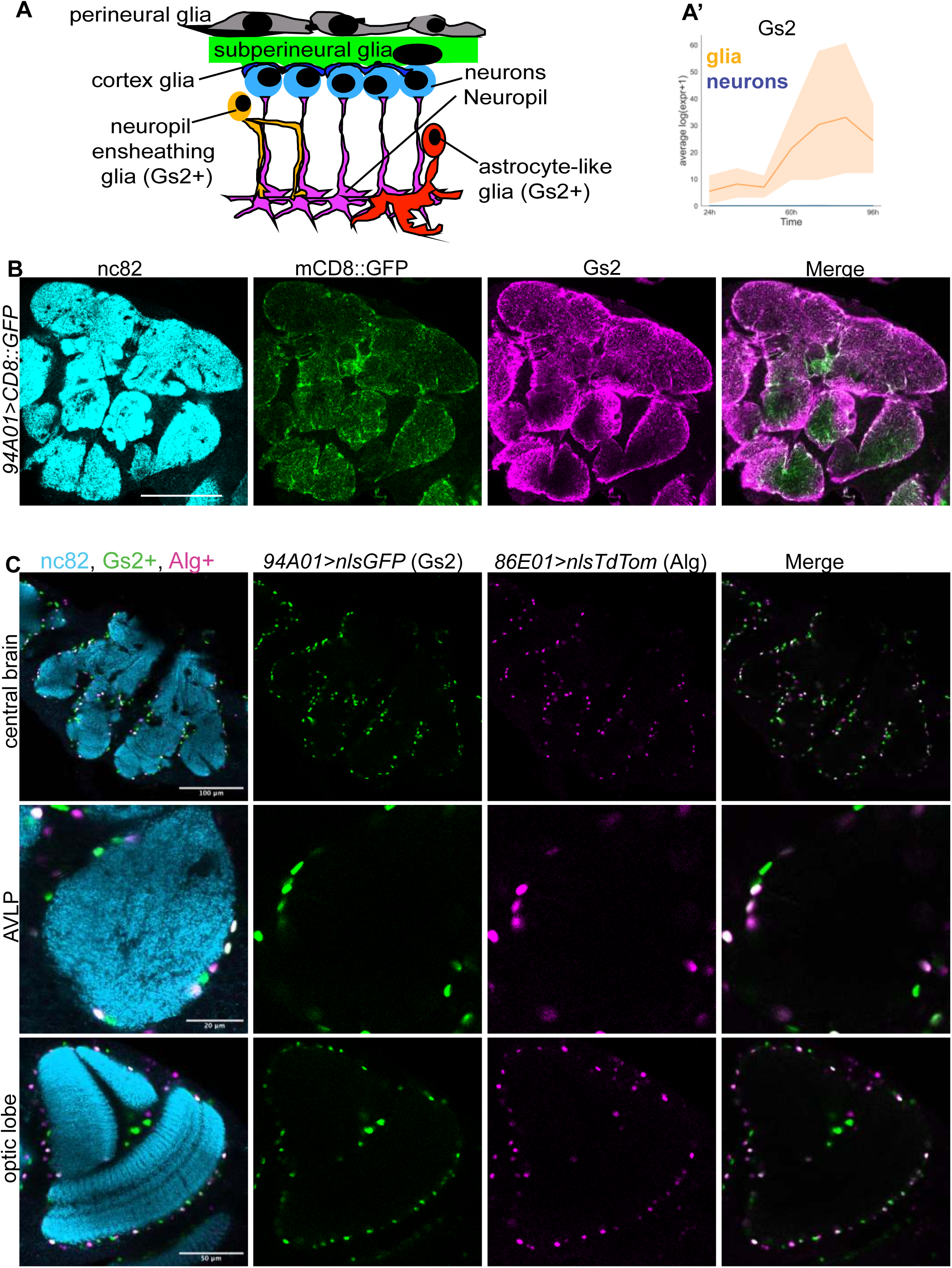
(A) Schematic of fly brain neurons and glia, with Gs2+ glia encompassing neuropil types composed of ensheathing (orange) and astrocyte-like (red) glia^85^. (A)’ Gs2 expression in scRNAseq datasets from^74^, with glia (orange) exclusively expressing Gs2, with expression increasing from 48h APF. (B) Gs2 enhancer trap (*GMR94A01*-*GAL4*) expressing *UAS-CD8::GFP* (green), stained with the neuropil marker nc82 (light blue) and Gs2 (magenta) in day 1 brains. Gs2+ GFP-labeled membranes are decorated by Gs2; note the absence of cortex and barrier signal. Scale bar = 100µm. n = 5 brains. (C) Day 1 adult brains labelingr Gs2+ cells with *GMR94A01-GAL4 >UAS-nls-STINGER* (green) and astrocyte-like glia (ALG) labeled with *GMR86E01-LexA > LexAop-nls-TdTom* (magenta) in central brain (scale bar = 100µm), AVLP (scale bar = 20µm), and optic lobe (scale bar = 50µm). Neuropil is stained with nc82 (light blue). n = 3 brains.

**Figure S5, related to Figure 5.**
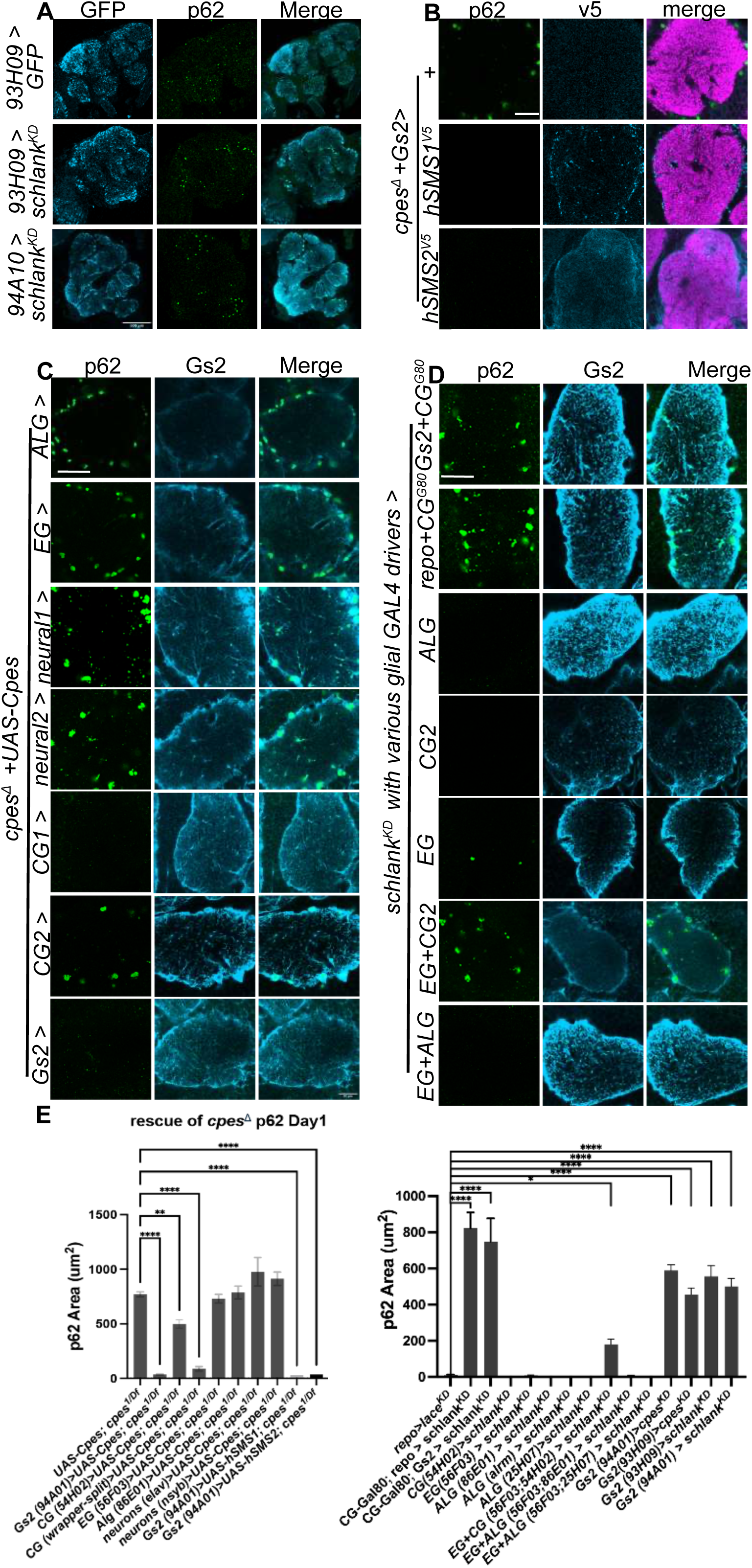
(A) Removing *schlank* (Ceramide Synthase) in Gs2 glia with two independent drivers (*GMR94A01-GAL4* or *GMR93H09-GAL4*) caused autonomous p62 accumulation (green) in GFP-labeled Gs2 membranes (light blue). Scale bar = 100 µm. (B) Expressing *hSMS1::v5* or *hSMS2::v5* in Gs2+ glia with *GMR94A01-GAL4* in *cpes* nulls rescued p62 (green). V5 (blue) was expressed in a pattern consistent with the predicted subcellular compartment of these enzymes (golgi for SMS1, which appears punctate; plasma membrane for SMS2). Magenta shows nc82 neuropil in the AVLP. Scale bar = 20 µm. (C) Glial or neuronal *GAL4* drivers re-expressing *UAS-Cpes* in attempted rescue of the *cpes* null mutant accumulation of p62 (green) in Gs2+ glia (light blue) in the AVLP. ALG = astrocyte-like glia (*GMR86E01-GAL4*); EG = ensheathing glia *(GMR56F03-GAL4*); neural1 = *elav-GAL4*; neural2 = *nSyb-GAL4*; CG1 = cortex glia (*cortex-split-GAL4*); CG2 = cortex glia (*GMR54H02-GAL4*); Gs2 = *94A01-GAL4*. Notably, Gs2 drivers fully rescued the p62 phenotype, while cortex glia drivers either fully rescued (*ctx-split*) or partially rescued (*GMR54H02*) p62 levels. Compensatory interactions between cortex glia and neuropil glia have been observed recently^154^. Scale bar = 20 µm. (D) Staining for p62 (green) in Gs2+ glia (light blue) by glial driver combinations crossed to *schlank-RNAi*, including *Gs2-GAL4* or *repo-GAL4* with GAL80 produced in cortex glia (*ctx^G^*^80^) to exclude leaky GAL4 expression in cortex glia. Outside of *Gs2-GAL4* or *repo-GAL4*, only the combined ensheathing glia + cortex glia knockdown (EG+CG > *schlank^KD^*using *GMR56F03-GAL4; GMR54H02-GAL4*) caused a partial p62 phenotype. Scale bar = 20 µm. (E) Quantification of p62 aggregates in AVLP from genotypes in A-D. Data are represented as mean ± SEM. n > 10 brains per condition. * p < 0.05, ** p < 0.01, *** p < 0.001, **** p < 0.0001

**Figure S6, related to Figure 6.**
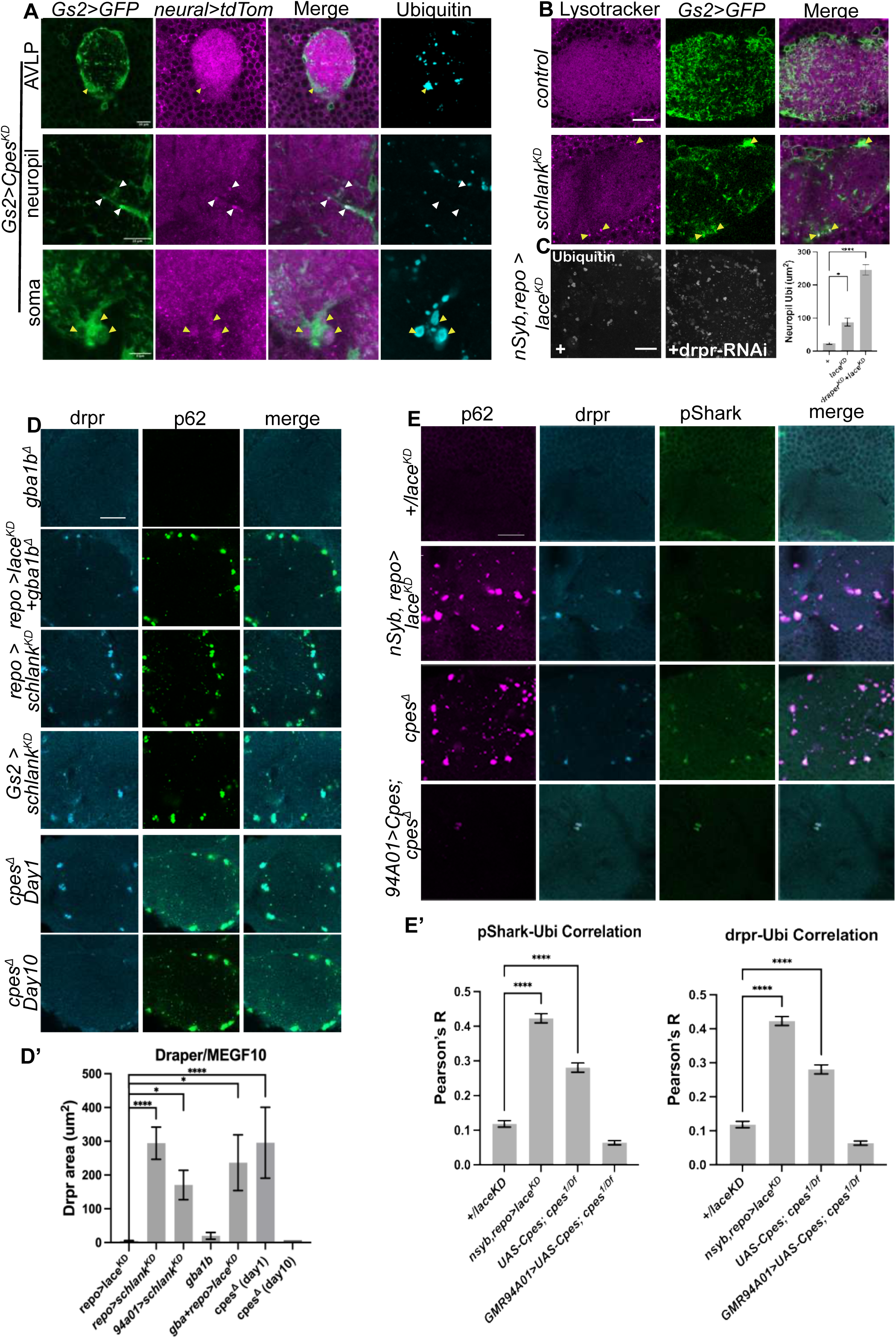
(A) Gs2 (green) and neuronal membranes (magenta) in the AVLP stained with ubiquitin (blue) in Gs2-glial knockdowns of *cpes* (scale bar = 20 µm), with zooms into neuropil (scale bar = 10 µm), and soma (scale bar = 5 µm). White arrows mark ubiquitin-, neuronal+ inclusions in glial membranes. Yellow arrows indicate ubiquitin+, weakly neuronal+ inclusions. (B) Lysotracker staining of Gs2 glial membranes in AVLP in controls and *schlank* knockdowns, with smaller lysotracker signals evident in Gs2 soma (arrows). Scale bar = 20 µm. (C) Removing *draper* worsens ubiquitin accumulation in brains with *lace* removed in both neurons and glia using *nSyb-GAL4, repo-GAL4*. Scale bar = 20 µm. (D-D’) Draper (light blue) accumulates in day 1 AVLP brains from genotypes that blockade CPE biosynthesis. Note that draper is lost from these structures by day 10 in *cpes* nulls. D’, quantification of draper accumulation. Scale bar = 20 µm. (E-E’) pShark staining in AVLP correlates with draper and p62 in genotypes with a blockade in CPE biosynthesis. E’, quantification. Scale bar = 20 µm. Data are represented as mean ± SEM. n > 7 brains per condition. * p < 0.05, ** p < 0.01, *** p < 0.001, **** p < 0.0001

**Figure S7, related to Figure 7.**
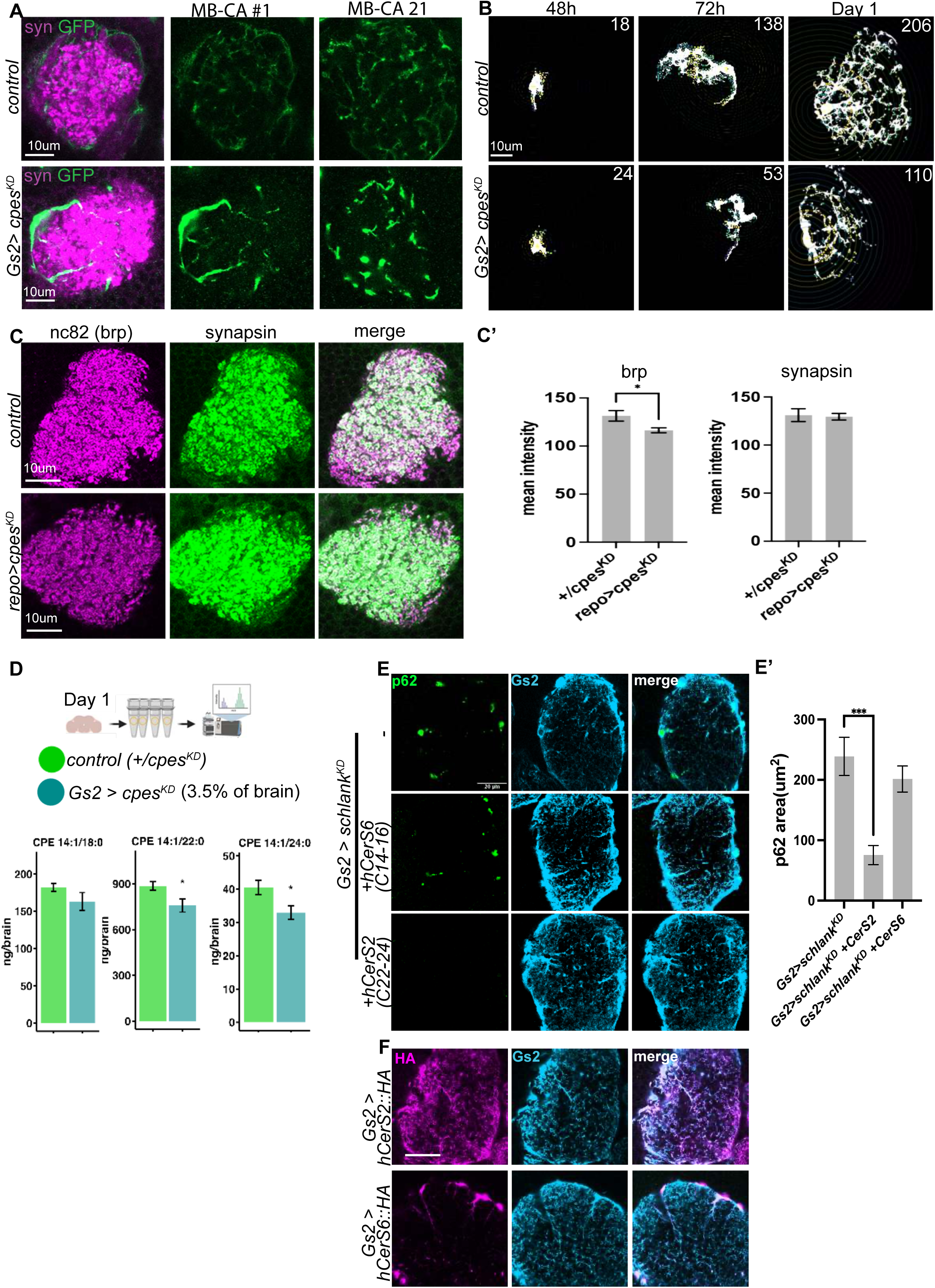
(A) Non-sparsely labeled Gs2 glial membranes (green) in control and *cpes-RNAi* mushroom body calyx (MB-CA). Syn = synapsin, neuropil marker in magenta. Aberrant thicker branches are observed in *cpes* knockdowns with *Gs2-GAL4*. Scale bars = 10 µm. (B) Example Sholl analysis of sparsely labeled glial clones from Figure 7. Scale bar = 10 µm. Numbers in top right indicate number of intersections, a proxy for branching. (C-C’) Mushroom body calyx stained for brp (nc82, magenta) or synapsin (green) from controls or glia depleted of *cpes*. Brains were pooled and stained in the same primary and secondary well, then decoded for the presence of *cpes-RNAi* by p62 aggregates. Brp/nc82 levels were reduced, unlike synapsin. Scale bar = 10 µm. (D) Lipidomics of controls (*+/cpes^KD^*, green) versus *cpes* knockdown in Gs2+ glia (blue) from day 1 brains. CPE 14:1/22:0 and CPE 14:1/24:0 are decreased, despite the genetic manipulation only targetting only ∼3.5% of the brain. (E) Rescue of p62 (green) in *schlank/CerS* knockdown AVLPs in Gs2 glia (light blue) by heterologous expression of human hCerS2 or hCerS6. CerS2 generates C22-C24 VLCFA sphingolipids, while CerS6 generates C14-C16 sphingolipids. Scale bar = 20 µm. (E’) Quantification of p62 area. (F) Localization of hCerS transgenes (magneta) by anti-HA staining when expressed in Gs2 glia (light blue). Scale bar = 20 µm. Data are represented as mean ± SEM. n > 10 brains per genotype for A-C and E-F, and 4 tubes of 15 brains each for D. * p < 0.05, ** p < 0.01, *** p < 0.001, **** p < 0.0001

## References

1. Südhof, T.C. (2018). Towards an Understanding of Synapse Formation. Neuron 100, 276–293. 10.1016/j.neuron.2018.09.040.

2. Luo, L., and O’Leary, D.D.M. (2005). AXON RETRACTION AND DEGENERATION IN DEVELOPMENT AND DISEASE. Annu. Rev. Neurosci. 28, 127–156. 10.1146/annurev.neuro.28.061604.135632.

3. Sanes, J.R., and Zipursky, S.L. (2020). Synaptic Specificity, Recognition Molecules, and Assembly of Neural Circuits. Cell 181, 536–556. 10.1016/j.cell.2020.04.008.

4. Holguera, I., and Desplan, C. (2018). Neuronal specification in space and time. Science362, 176–180. 10.1126/science.aas9435.

5. Garcia, M.A., and Zuchero, J.B. (2019). Anchors Away: Glia-Neuron Adhesion Regulates Myelin Targeting and Growth. Dev. Cell 51, 659–661. 10.1016/j.devcel.2019.11.018.

6. Allen, N.J., and Lyons, D.A. (2018). Glia as architects of central nervous system formation and function. Science 362, 181–185. 10.1126/science.aat0473.

7. Barres, B.A. (2008). The Mystery and Magic of Glia: A Perspective on Their Roles in Health and Disease. Neuron 60, 430–440. 10.1016/j.neuron.2008.10.013.

8. Hartenstein, V. (2011). Morphological diversity and development of glia in Drosophila. Glia 59, 1237–1252. 10.1002/glia.21162.

9. Yildirim, K., Petri, J., Kottmeier, R., and Klämbt, C. (2019). Drosophila glia: Few cell types and many conserved functions. Glia 67, 5–26. 10.1002/glia.23459.

10. Schwabe, T., Neuert, H., and Clandinin, T.R. (2013). A Network of Cadherin-Mediated Interactions Polarizes Growth Cones to Determine Targeting Specificity. Cell 154, 351–364. 10.1016/j.cell.2013.06.011.

11. Brockschnieder, D., Sabanay, H., Riethmacher, D., and Peles, E. (2006). Ermin, A Myelinating Oligodendrocyte-Specific Protein That Regulates Cell Morphology. J. Neurosci. 26, 757–762. 10.1523/JNEUROSCI.4317-05.2006.

12. Zuchero, J.B., Fu, M., Sloan, S.A., Ibrahim, A., Olson, A., Zaremba, A., Dugas, J.C., Wienbar, S., Caprariello, A.V., Kantor, C., et al. (2015). CNS Myelin Wrapping Is Driven by Actin Disassembly. Dev. Cell 34, 152–167. 10.1016/j.devcel.2015.06.011.

13. Xu, C., Theisen, E., Maloney, R., Peng, J., Santiago, I., Yapp, C., Werkhoven, Z., Rumbaut, E., Shum, B., Tarnogorska, D., et al. (2019). Control of Synaptic Specificity by Establishing a Relative Preference for Synaptic Partners. Neuron 103, 865–877.e7. 10.1016/j.neuron.2019.06.006.

14. McLaughlin, C.N., Perry-Richardson, J.J., Coutinho-Budd, J.C., and Broihier, H.T. (2019). Dying Neurons Utilize Innate Immune Signaling to Prime Glia for Phagocytosis during Development. Dev. Cell 48, 506–522.e6. 10.1016/j.devcel.2018.12.019.

15. Hartfuss, E., Förster, E., Bock, H.H., Hack, M.A., Leprince, P., Luque, J.M., Herz, J., Frotscher, M., and Götz, M. (2003). Reelin signaling directly affects radial glia morphology and biochemical maturation. Development 130, 4597–4609. 10.1242/dev.00654.

16. Stogsdill, J.A., Ramirez, J., Liu, D., Kim, Y.H., Baldwin, K.T., Enustun, E., Ejikeme, T., Ji, R.-R., and Eroglu, C. (2017). Astrocytic neuroligins control astrocyte morphogenesis and synaptogenesis. Nature 551, 192–197. 10.1038/nature24638.

17. Lam, M., Takeo, K., Almeida, R.G., Cooper, M.H., Wu, K., Iyer, M., Kantarci, H., and Zuchero, J.B. (2022). CNS myelination requires VAMP2/3-mediated membrane expansion in oligodendrocytes. Nat. Commun. 13, 5583. 10.1038/s41467-022-33200-4.

18. Freeman, M.R. (2010). Specification and Morphogenesis of Astrocytes. Science 330, 774–778. 10.1126/science.1190928.

19. Eroglu, C., and Barres, B.A. (2010). Regulation of synaptic connectivity by glia. Nature 468, 223–231. 10.1038/nature09612.

20. Allen, N.J., and Eroglu, C. (2017). Cell Biology of Astrocyte-Synapse Interactions. Neuron 96, 697–708. 10.1016/j.neuron.2017.09.056.

21. Wallroth, A., and Haucke, V. (2018). Phosphoinositide conversion in endocytosis and the endolysosomal system. J. Biol. Chem. 293, 1526–1535. 10.1074/jbc.R117.000629.

22. Wolf, S.A., Boddeke, H.W.G.M., and Kettenmann, H. (2017). Microglia in Physiology and Disease. Annu. Rev. Physiol. 79, 619–643. 10.1146/annurev-physiol-022516-034406.

23. Watts, R.J., Schuldiner, O., Perrino, J., Larsen, C., and Luo, L. (2004). Glia Engulf Degenerating Axons during Developmental Axon Pruning. Curr. Biol. 14, 678–684. 10.1016/j.cub.2004.03.035.

24. Awasaki, T., and Ito, K. (2004). Engulfing action of glial cells is required for programmed axon pruning during Drosophila metamorphosis. Curr. Biol. CB 14, 668–677. 10.1016/j.cub.2004.04.001.

25. Doherty, J., Logan, M.A., Tasdemir, O.E., and Freeman, M.R. (2009). Ensheathing Glia Function as Phagocytes in the Adult Drosophila Brain. J. Neurosci. 29, 4768–4781. 10.1523/JNEUROSCI.5951-08.2009.

26. Roney, J.C., Cheng, X.-T., and Sheng, Z.-H. (2022). Neuronal endolysosomal transport and lysosomal functionality in maintaining axonostasis. J. Cell Biol. 221, e202111077. 10.1083/jcb.202111077.

27. Baba, T., Toth, D.J., Sengupta, N., Kim, Y.J., and Balla, T. (2019). Phosphatidylinositol 4,5-bisphosphate controls Rab7 and PLEKHM1 membrane cycling during autophagosome–lysosome fusion. EMBO J. 38, e100312. 10.15252/embj.2018100312.

28. Koike, S., and Jahn, R. (2024). Rab GTPases and phosphoinositides fine-tune SNAREs dependent targeting specificity of intracellular vesicle traffic. Nat. Commun. 15, 2508. 10.1038/s41467-024-46678-x.

29. Kolter, T., and Sandhoff, K. (2005). PRINCIPLES OF LYSOSOMAL MEMBRANE DIGESTION: Stimulation of Sphingolipid Degradation by Sphingolipid Activator Proteins and Anionic Lysosomal Lipids. Annu. Rev. Cell Dev. Biol. 21, 81–103. 10.1146/annurev.cellbio.21.122303.120013.

30. Nyame, K., Hims, A., Aburous, A., Laqtom, N.N., Dong, W., Medoh, U.N., Heiby, J.C., Xiong, J., Ori, A., and Abu-Remaileh, M. (2024). Glycerophosphodiesters inhibit lysosomal phospholipid catabolism in Batten disease. Mol. Cell 84, 1354–1364.e9. 10.1016/j.molcel.2024.02.006.

31. Shayman, J.A., and Tesmer, J.J.G. (2019). Lysosomal phospholipase A2. Biochim. Biophys. Acta BBA - Mol. Cell Biol. Lipids 1864, 932–940. 10.1016/j.bbalip.2018.07.012.

32. Svennerholm, L. (1968). Distribution and fatty acid composition of phosphoglycerides in normal human brain. J. Lipid Res. 9, 570–579. 10.1016/S0022-2275(20)42702-6.

33. Svennerholm, L., Boström, K., Jungbjer, B., and Olsson, L. (1994). Membrane Lipids of Adult Human Brain: Lipid Composition of Frontal and Temporal Lobe in Subjects of Age 20 to 100 Years. J. Neurochem. 63, 1802–1811. 10.1046/j.1471-4159.1994.63051802.x.

34. Fitzner, D., Bader, J.M., Penkert, H., Bergner, C.G., Su, M., Weil, M.-T., Surma, M.A., Mann, M., Klose, C., and Simons, M. (2020). Cell-Type- and Brain-Region-Resolved Mouse Brain Lipidome. Cell Rep. 32, 108132. 10.1016/j.celrep.2020.108132.

35. O’Brien, J.S., and Sampson, E.L. (1965). Fatty acid and fatty aldehyde composition of the major brain lipids in normal human gray matter, white matter, and myelin. J. Lipid Res. 6, 545–551.

36. Bozek, K., Wei, Y., Yan, Z., Liu, X., Xiong, J., Sugimoto, M., Tomita, M., Pääbo, S., Sherwood, C.C., Hof, P.R., et al. (2015). Organization and Evolution of Brain Lipidome Revealed by Large-Scale Analysis of Human, Chimpanzee, Macaque, and Mouse Tissues. Neuron 85, 695–702. 10.1016/j.neuron.2015.01.003.

37. Piomelli, D., Astarita, G., and Rapaka, R. (2007). A neuroscientist’s guide to lipidomics. Nat. Rev. Neurosci. 8, 743–754. 10.1038/nrn2233.

38. Levental, I., and Lyman, E. (2022). Regulation of membrane protein structure and function by their lipid nano-environment. Nat. Rev. Mol. Cell Biol., 1–16. 10.1038/s41580-022-00524-4.

39. Holthuis, J.C.M., and Levine, T.P. (2005). Lipid traffic: floppy drives and a superhighway. Nat. Rev. Mol. Cell Biol. 6, 209–220. 10.1038/nrm1591.

40. Naruse, H., Ishiura, H., Esaki, K., Mitsui, J., Satake, W., Greimel, P., Shingai, N., Machino, Y., Kokubo, Y., Hamaguchi, H., et al. (2024). variants are associated with early-onset ALS and FTD due to aberrant sphingolipid synthesis. Ann. Clin. Transl. Neurol. 11, 946–957. 10.1002/acn3.52013.

41. Lin, G., Wang, L., Marcogliese, P.C., and Bellen, H.J. (2019). Sphingolipids in the Pathogenesis of Parkinson’s Disease and Parkinsonism. Trends Endocrinol. Metab. 30, 106–117. 10.1016/j.tem.2018.11.003.

42. Sidransky, E., and Lopez, G. (2012). The link between the GBA gene and parkinsonism. Lancet Neurol. 11, 986–998. 10.1016/S1474-4422(12)70190-4.

43. Grimm, M.O.W., Grimm, H.S., Pätzold, A.J., Zinser, E.G., Halonen, R., Duering, M., Tschäpe, J.-A., Strooper, B.D., Müller, U., Shen, J., et al. (2005). Regulation of cholesterol and sphingomyelin metabolism by amyloid-β and presenilin. Nat. Cell Biol. 7, 1118– 1123. 10.1038/ncb1313.

44. Ariga, T., McDonald, M.P., and Yu, R.K. (2008). Thematic Review Series: Sphingolipids. Role of ganglioside metabolism in the pathogenesis of Alzheimer’s disease—a review,. J. Lipid Res. 49, 1157–1175. 10.1194/jlr.R800007-JLR200.

45. Bandaru, V.V.R., Troncoso, J., Wheeler, D., Pletnikova, O., Wang, J., Conant, K., and Haughey, N.J. (2009). ApoE4 disrupts sterol and sphingolipid metabolism in Alzheimer’s but not normal brain. Neurobiol. Aging 30, 591–599. 10.1016/j.neurobiolaging.2007.07.024.

46. Mohassel, P., Donkervoort, S., Lone, M.A., Nalls, M., Gable, K., Gupta, S.D., Foley, A.R., Hu, Y., Saute, J.A.M., Moreira, A.L., et al. (2021). Childhood amyotrophic lateral sclerosis caused by excess sphingolipid synthesis. Nat. Med. 27, 1197–1204. 10.1038/s41591-021-01346-1.

47. Furuya, S., Mitoma, J., Makino, A., and Hirabayashi, Y. (1998). Ceramide and Its Interconvertible Metabolite Sphingosine Function as Indispensable Lipid Factors Involved in Survival and Dendritic Differentiation of Cerebellar Purkinje Cells. J. Neurochem. 71, 366–377. 10.1046/j.1471-4159.1998.71010366.x.

48. Schwarz, A., and Futerman, A.H. (1997). Distinct Roles for Ceramide and Glucosylceramide at Different Stages of Neuronal Growth. J. Neurosci. 17, 2929–2938. 10.1523/JNEUROSCI.17-09-02929.1997.

49. Barnes-Vélez, J.A., Yasar, F.B.A., and Hu, J. (2023). Myelin lipid metabolism and its role in myelination and myelin maintenance. The Innovation 4. 10.1016/j.xinn.2022.100360.

50. Coetzee, T., Fujita, N., Dupree, J., Shi, R., Blight, A., Suzuki, K., Suzuki, K., and Popko, B. (1996). Myelination in the absence of galactocerebroside and sulfatide: normal structure with abnormal function and regional instability. Cell 86, 209–219. 10.1016/s0092-8674(00)80093-8.

51. Marcus, J., Honigbaum, S., Shroff, S., Honke, K., Rosenbluth, J., and Dupree, J. l. (2006). Sulfatide is essential for the maintenance of CNS myelin and axon structure. Glia 53, 372–381. 10.1002/glia.20292.

52. Imgrund, S., Hartmann, D., Farwanah, H., Eckhardt, M., Sandhoff, R., Degen, J., Gieselmann, V., Sandhoff, K., and Willecke, K. (2009). Adult Ceramide Synthase 2 (CERS2)-deficient Mice Exhibit Myelin Sheath Defects, Cerebellar Degeneration, and Hepatocarcinomas *. J. Biol. Chem. 284, 33549–33560. 10.1074/jbc.M109.031971.

53. Liu, H., Wang, X., Chen, L., Chen, L., Tsirka, S.E., Ge, S., and Xiong, Q. (2021). Microglia modulate stable wakefulness via the thalamic reticular nucleus in mice. Nat. Commun. 12, 4646. 10.1038/s41467-021-24915-x.

54. Flury, A., Aljayousi, L., Park, H.-J., Khakpour, M., Mechler, J., Aziz, S., McGrath, J.D., Deme, P., Sandberg, C., Ibáñez, F.G., et al. (2025). A neurodegenerative cellular stress response linked to dark microglia and toxic lipid secretion. Neuron 113, 554–571.e14. 10.1016/j.neuron.2024.11.018.

55. Fan, Y., Soller, M., Flister, S., Hollmann, M., Müller, M., Bello, B., Egger, B., White, K., Schäfer, M.A., and Reichert, H. (2005). The egghead gene is required for compartmentalization in Drosophila optic lobe development. Dev. Biol. 287, 61–73. 10.1016/j.ydbio.2005.08.031.

56. Goyal, G., Zheng, J., Adam, E., Steffes, G., Jain, M., Klavins, K., and Hummel, T. (2019). Sphingolipid-dependent Dscam sorting regulates axon segregation. Nat. Commun. 10, 813. 10.1038/s41467-019-08765-2.

57. Midorikawa, R., Yamamoto-Hino, M., Awano, W., Hinohara, Y., Suzuki, E., Ueda, R., and Goto, S. (2010). Autophagy-Dependent Rhodopsin Degradation Prevents Retinal Degeneration in Drosophila. J. Neurosci. 30, 10703–10719. 10.1523/JNEUROSCI.2061-10.2010.

58. Hebbar, S., Schuhmann, K., Shevchenko, A., and Knust, E. (2020). Hydroxylated sphingolipid biosynthesis regulates photoreceptor apical domain morphogenesis. J. Cell Biol. 219, e201911100. 10.1083/jcb.201911100.

59. Kunduri, G., Turner-Evans, D., Konya, Y., Izumi, Y., Nagashima, K., Lockett, S., Holthuis, J., Bamba, T., Acharya, U., and Acharya, J.K. (2018). Defective cortex glia plasma membrane structure underlies light-induced epilepsy in cpes mutants. Proc. Natl. Acad. Sci. U. S. A. 115, E8919–E8928. 10.1073/pnas.1808463115.

60. Ghosh, A., Kling, T., Snaidero, N., Sampaio, J.L., Shevchenko, A., Gras, H., Geurten, B., Göpfert, M.C., Schulz, J.B., Voigt, A., et al. (2013). A Global In Vivo Drosophila RNAi Screen Identifies a Key Role of Ceramide Phosphoethanolamine for Glial Ensheathment of Axons. PLOS Genet. 9, e1003980. 10.1371/journal.pgen.1003980.

61. Kottmeier, R., Bittern, J., Schoofs, A., Scheiwe, F., Matzat, T., Pankratz, M., and Klämbt, C. (2020). Wrapping glia regulates neuronal signaling speed and precision in the peripheral nervous system of Drosophila. Nat. Commun. 11, 4491. 10.1038/s41467-020-18291-1.

62. Chung, H., Ye, Q., Park, Y.-J., Zuo, Z., Mok, J.-W., Kanca, O., Tattikota, S.G., Lu, S., Perrimon, N., Lee, H.K., et al. (2023). Very long-chain fatty acids induce glial-derived Sphingosine-1-Phosphate synthesis, secretion, and neuroinflammation. Cell Metab. 35, 855–874.e5. 10.1016/j.cmet.2023.03.022.

63. Kinghorn, K.J., Grönke, S., Castillo-Quan, J.I., Woodling, N.S., Li, L., Sirka, E., Gegg, M., Mills, K., Hardy, J., Bjedov, I., et al. (2016). A Drosophila Model of Neuronopathic Gaucher Disease Demonstrates Lysosomal-Autophagic Defects and Altered mTOR Signalling and Is Functionally Rescued by Rapamycin. J. Neurosci. Ok. J. Soc. Neurosci. 36, 11654–11670. 10.1523/JNEUROSCI.4527-15.2016.

64. Davis, M.Y., Trinh, K., Thomas, R.E., Yu, S., Germanos, A.A., Whitley, B.N., Sardi, S.P., Montine, T.J., and Pallanck, L.J. (2016). Glucocerebrosidase Deficiency in Drosophila Results in α-Synuclein-Independent Protein Aggregation and Neurodegeneration. PLOS Genet. 12, e1005944. 10.1371/journal.pgen.1005944.

65. Vaughen, J.P., Theisen, E., Rivas-Serna, I.M., Berger, A.B., Kalakuntla, P., Anreiter, I., Mazurak, V.C., Rodriguez, T.P., Mast, J.D., Hartl, T., et al. (2022). Glial control of sphingolipid levels sculpts diurnal remodeling in a circadian circuit. Neuron 110, 3186–3205.e7. 10.1016/j.neuron.2022.07.016.

66. Hindle, S.J., Hebbar, S., Schwudke, D., Elliott, C.J.H., and Sweeney, S.T. (2017). A saposin deficiency model in Drosophila: Lysosomal storage, progressive neurodegeneration and sensory physiological decline. Neurobiol. Dis. 98, 77–87. 10.1016/j.nbd.2016.11.012.

67. Sellin, J., Schulze, H., Paradis, M., Gosejacob, D., Papan, C., Shevchenko, A., Psathaki, O.E., Paululat, A., Thielisch, M., Sandhoff, K., et al. (2017). Characterization of Drosophila Saposin-related mutants as a model for lysosomal sphingolipid storage diseases. Dis. Model. Mech. 10, 737–750. 10.1242/dmm.027953.

68. Hebbar, S., Khandelwal, A., Jayashree, R., Hindle, S.J., Chiang, Y.N., Yew, J.Y., Sweeney, S.T., and Schwudke, D. (2017). Lipid metabolic perturbation is an early-onset phenotype in adult spinster mutants: a Drosophila model for lysosomal storage disorders. Mol. Biol. Cell 28, 3728–3740. 10.1091/mbc.e16-09-0674.

69. Chen, X., Li, J., Gao, Z., Yang, Y., Kuang, W., Dong, Y., Chua, G.H., Huang, X., Jiang, B., Tian, H., et al. (2022). Endogenous ceramide phosphoethanolamine modulates circadian rhythm via neural–glial coupling in *Drosophila*. Natl. Sci. Rev. 9, nwac148. 10.1093/nsr/nwac148.

70. Wang, L., Lin, G., Zuo, Z., Li, Y., Byeon, S.K., Pandey, A., and Bellen, H.J. (2022). Neuronal activity induces glucosylceramide that is secreted via exosomes for lysosomal degradation in glia. Sci. Adv. 8, eabn3326. 10.1126/sciadv.abn3326.

71. Cabasso, O., Paul, S., Dorot, O., Maor, G., Krivoruk, O., Pasmanik-Chor, M., Mirzaian, M., Ferraz, M., Aerts, J., and Horowitz, M. (2019). Drosophila melanogaster Mutated in its GBA1b Ortholog Recapitulates Neuronopathic Gaucher Disease. J. Clin. Med. 8, 1420. 10.3390/jcm8091420.

72. Akin, O., Bajar, B.T., Keles, M.F., Frye, M.A., and Zipursky, S.L. (2019). Cell-type-Specific Patterned Stimulus-Independent Neuronal Activity in the Drosophila Visual System during Synapse Formation. Neuron 101, 894–904.e5. 10.1016/j.neuron.2019.01.008.

73. Muthukumar, A.K., Stork, T., and Freeman, M.R. (2014). Activity-dependent regulation of astrocyte GAT levels during synaptogenesis. Nat. Neurosci. 17, 1340–1350. 10.1038/nn.3791.

74. Kurmangaliyev, Y.Z., Yoo, J., Valdes-Aleman, J., Sanfilippo, P., and Zipursky, S.L. (2020). Transcriptional Programs of Circuit Assembly in the Drosophila Visual System. Neuron 108, 1045–1057.e6. 10.1016/j.neuron.2020.10.006.

75. Leier, H.C., Foden, A.J., Jindal, D.A., Wilkov, A.J., Costello, P.V. der L., Vanderzalm, P.J., Coutinho-Budd, J.C., Tabuchi, M., and Broihier, H.T. (2025). Glia control experience-dependent plasticity in an olfactory critical period. eLife 13. 10.7554/eLife.100989.2.

76. Carvalho, M., Schwudke, D., Sampaio, J.L., Palm, W., Riezman, I., Dey, G., Gupta, G.D., Mayor, S., Riezman, H., Shevchenko, A., et al. (2010). Survival strategies of a sterol auxotroph. Dev. Camb. Engl. 137, 3675–3685. 10.1242/dev.044560.

77. Carvalho, M., Sampaio, J.L., Palm, W., Brankatschk, M., Eaton, S., and Shevchenko, A. (2012). Effects of diet and development on the Drosophila lipidome. Mol. Syst. Biol. 8, 600. 10.1038/msb.2012.29.

78. Kumar, M., Has, C., Lam-Kamath, K., Ayciriex, S., Dewett, D., Bashir, M., Poupault, C., Schuhmann, K., Thomas, H., Knittelfelder, O., et al. (2024). Lipidome Unsaturation Affects the Morphology and Proteome of the Drosophila Eye. J. Proteome Res. 23, 1188– 1199. 10.1021/acs.jproteome.3c00570.

79. Lee, P.-T., Zirin, J., Kanca, O., Lin, W.-W., Schulze, K.L., Li-Kroeger, D., Tao, R., Devereaux, C., Hu, Y., Chung, V., et al. (2018). A gene-specific T2A-GAL4 library for Drosophila. eLife 7, e35574. 10.7554/eLife.35574.

80. Fyrst, H., Herr, D.R., Harris, G.L., and Saba, J.D. (2004). Characterization of free endogenous C14 and C16 sphingoid bases from Drosophila melanogaster. J. Lipid Res. 45, 54–62. 10.1194/jlr.M300005-JLR200.

81. Hull, A.J., Atilano, M.L., Hallqvist, J., Heywood, W., and Kinghorn, K.J. (2024). Ceramide lowering rescues respiratory defects in a Drosophila model of acid sphingomyelinase deficiency. Hum. Mol. Genet. 33, 2111–2122. 10.1093/hmg/ddae143.

82. Jewett, K.A., Thomas, R.E., Phan, C.Q., Lin, B., Milstein, G., Yu, S., Bettcher, L.F., Neto, F.C., Djukovic, D., Raftery, D., et al. (2021). Glucocerebrosidase reduces the spread of protein aggregation in a Drosophila melanogaster model of neurodegeneration by regulating proteins trafficked by extracellular vesicles. PLoS Genet. 17, e1008859. 10.1371/journal.pgen.1008859.

83. Katsuragi, Y., Ichimura, Y., and Komatsu, M. (2015). p62/SQSTM1 functions as a signaling hub and an autophagy adaptor. FEBS J. 282, 4672–4678. 10.1111/febs.13540.

84. Kremer, M.C., Jung, C., Batelli, S., Rubin, G.M., and Gaul, U. (2017). The glia of the adult *Drosophila* nervous system: Glia Anatomy in Adult Drosophila Nervous System. Glia 65, 606–638. 10.1002/glia.23115.

85. Kato, K., Orihara-Ono, M., and Awasaki, T. (2020). Multiple lineages enable robust development of the neuropil-glia architecture in adult *Drosophila*. Development 147, dev184085. 10.1242/dev.184085.

86. Vacaru, A.M., Van Den Dikkenberg, J., Ternes, P., and Holthuis, J.C.M. (2013). Ceramide Phosphoethanolamine Biosynthesis in Drosophila Is Mediated by a Unique Ethanolamine Phosphotransferase in the Golgi Lumen. J. Biol. Chem. 288, 11520–11530. 10.1074/jbc.M113.460972.

87. Davis, D.L., Mahawar, U., Pope, V.S., Allegood, J., Sato-Bigbee, C., and Wattenberg, B.W. (2020). Dynamics of sphingolipids and the serine palmitoyltransferase complex in rat oligodendrocytes during myelination [S]. J. Lipid Res. 61, 505–522. 10.1194/jlr.RA120000627.

88. Bhat, H.B., Ishitsuka, R., Inaba, T., Murate, M., Abe, M., Makino, A., Kohyama-Koganeya, A., Nagao, K., Kurahashi, A., Kishimoto, T., et al. (2015). Evaluation of aegerolysins as novel tools to detect and visualize ceramide phosphoethanolamine, a major sphingolipid in invertebrates. FASEB J. 29, 3920–3934. 10.1096/fj.15-272112.

89. Kunduri, G., Le, S.-H., Baena, V., Vijaykrishna, N., Harned, A., Nagashima, K., Blankenberg, D., Yoshihiro, I., Narayan, K., Bamba, T., et al. (2022). Delivery of ceramide phosphoethanolamine lipids to the cleavage furrow through the endocytic pathway is essential for male meiotic cytokinesis. PLOS Biol. 20, e3001599. 10.1371/journal.pbio.3001599.

90. Bhat, H.B., Kishimoto, T., Abe, M., Makino, A., Inaba, T., Murate, M., Dohmae, N., Kurahashi, A., Nishibori, K., Fujimori, F., et al. (2013). Binding of a pleurotolysin ortholog from Pleurotus eryngii to sphingomyelin and cholesterol-rich membrane domains [S]. J. Lipid Res. 54, 2933–2943. 10.1194/jlr.D041731.

91. Tomita, T., Noguchi, K., Mimuro, H., Ukaji, F., Ito, K., Sugawara-Tomita, N., and Hashimoto, Y. (2004). Pleurotolysin, a Novel Sphingomyelin-specific Two-component Cytolysin from the Edible Mushroom Pleurotus ostreatus, Assembles into a Transmembrane Pore Complex *. J. Biol. Chem. 279, 26975–26982. 10.1074/jbc.M402676200.

92. Bernheimer, A.W., and Avigad, L.S. (1979). A cytolytic protein from the edible mushroom, Pleurotus ostreatus. Biochim. Biophys. Acta BBA - Gen. Subj. 585, 451–461. 10.1016/0304-4165(79)90090-4.

93. Freeman, M.R., Delrow, J., Kim, J., Johnson, E., and Doe, C.Q. (2003). Unwrapping Glial Biology: Gcm Target Genes Regulating Glial Development, Diversification, and Function. Neuron 38, 567–580. 10.1016/S0896-6273(03)00289-7.

94. Boulanger, A., Thinat, C., Züchner, S., Fradkin, L.G., Lortat-Jacob, H., and Dura, J.-M. (2021). Axonal chemokine-like Orion induces astrocyte infiltration and engulfment during mushroom body neuronal remodeling. Nat. Commun. 12, 1849. 10.1038/s41467-021-22054-x.

95. Ziegenfuss, J.S., Biswas, R., Avery, M.A., Hong, K., Sheehan, A.E., Yeung, Y.-G., Stanley, E.R., and Freeman, M.R. (2008). Draper-dependent glial phagocytic activity is mediated by Src and Syk family kinase signalling. Nature 453, 935–939. 10.1038/nature06901.

96. Stork, T., Sheehan, A., Tasdemir-Yilmaz, O.E., and Freeman, M.R. (2014). Neuron-Glia Interactions through the Heartless FGF Receptor Signaling Pathway Mediate Morphogenesis of Drosophila Astrocytes. Neuron 83, 388–403. 10.1016/j.neuron.2014.06.026.

97. Isaacman-Beck, J., Paik, K.C., Wienecke, C.F.R., Yang, H.H., Fisher, Y.E., Wang, I.E., Ishida, I.G., Maimon, G., Wilson, R.I., and Clandinin, T.R. (2020). SPARC enables genetic manipulation of precise proportions of cells. Nat. Neurosci. 23, 1168–1175. 10.1038/s41593-020-0668-9.

98. Ullian, E.M., Sapperstein, S.K., Christopherson, K.S., and Barres, B.A. (2001). Control of Synapse Number by Glia. Science 291, 657–661. 10.1126/science.291.5504.657.

99. Jay, T.R., Kang, Y., Jefferson, A., and Freeman, M.R. (2021). An ELISA-based method for rapid genetic screens in Drosophila. Proc. Natl. Acad. Sci. 118, e2107427118. 10.1073/pnas.2107427118.

100. Jain, S., Lin, Y., Kurmangaliyev, Y.Z., Valdes-Aleman, J., LoCascio, S.A., Mirshahidi, P., Parrington, B., and Zipursky, S.L. (2022). A global timing mechanism regulates cell-type-specific wiring programmes. Nature 603, 112–118. 10.1038/s41586-022-04418-5.

101. Vaughen, J.P., Theisen, E., and Clandinin, T.R. (2023). From seconds to days: Neural plasticity viewed through a lipid lens. Curr. Opin. Neurobiol. 80, 102702. 10.1016/j.conb.2023.102702.

102. Faust, T.E., Gunner, G., and Schafer, D.P. (2021). Mechanisms governing activity-dependent synaptic pruning in the developing mammalian CNS. Nat. Rev. Neurosci. 22, 657–673. 10.1038/s41583-021-00507-y.

103. Nelson, N., Vita, D.J., and Broadie, K. (2024). Experience-dependent glial pruning of synaptic glomeruli during the critical period. Sci. Rep. 14, 9110. 10.1038/s41598-024-59942-3.

104. Jindal, D.A., Leier, H.C., Salazar, G., Foden, A.J., Seitz, E.A., Wilkov, A.J., Coutinho-Budd, J.C., and Broihier, H.T. (2023). Early Draper-mediated glial refinement of neuropil architecture and synapse number in the Drosophila antennal lobe. Front. Cell. Neurosci. 17. 10.3389/fncel.2023.1166199.

105. Hakim-Mishnaevski, K., Flint-Brodsly, N., Shklyar, B., Levy-Adam, F., and Kurant, E. (2019). Glial Phagocytic Receptors Promote Neuronal Loss in Adult Drosophila Brain. Cell Rep. 29, 1438–1448.e3. 10.1016/j.celrep.2019.09.086.

106. Torres, A.Y., Nano, M., Campanale, J.P., Deak, S., and Montell, D.J. (2023). Activated Src kinase promotes cell cannibalism in *Drosophila*. J. Cell Biol. 222, e202302076. 10.1083/jcb.202302076.

107. Mishra, A.K., Rodriguez, M., Torres, A.Y., Smith, M., Rodriguez, A., Bond, A., Morrissey, M.A., and Montell, D.J. (2023). Hyperactive Rac stimulates cannibalism of living target cells and enhances CAR-M-mediated cancer cell killing. Proc. Natl. Acad. Sci. U. S. A. 120, e2310221120. 10.1073/pnas.2310221120.

108. Lennartz, H.M., Nunes, S. da C., Böhlig, K., Kuhn, S.M., Moneta, L., Yau, W.Y., Opálka, L., Iglesias-Artola, J.M., Modes, C.D., Honigmann, A., et al. (2025). Quantification of lipid sorting during clathrin-mediated endocytosis. Preprint at bioRxiv, 10.1101/2025.03.04.641423 https://doi.org/10.1101/2025.03.04.641423.

109. Pattingre, S., Bauvy, C., Levade, T., Levine, B., and Codogno, P. (2009). Ceramide-induced autophagy: To junk or to protect cells? Autophagy 5, 558–560. 10.4161/auto.5.4.8390.

110. Wu, C.-Y., Jhang, J.-G., Lin, W.-S., Chuang, P.-H., Lin, C.-W., Chu, L.-A., Chiang, A.-S., Ho, H.-C., Chan, C.-C., and Huang, S.-Y. (2021). Dihydroceramide desaturase promotes the formation of intraluminal vesicles and inhibits autophagy to increase exosome production. iScience 24, 103437. 10.1016/j.isci.2021.103437.

111. Sandhoff, R., and Sandhoff, K. (2018). Emerging concepts of ganglioside metabolism. FEBS Lett. 592, 3835–3864. 10.1002/1873-3468.13114.

112. Gomez-Sanchez, J.A., Carty, L., Iruarrizaga-Lejarreta, M., Palomo-Irigoyen, M., Varela-Rey, M., Griffith, M., Hantke, J., Macias-Camara, N., Azkargorta, M., Aurrekoetxea, I., et al. (2015). Schwann cell autophagy, myelinophagy, initiates myelin clearance from injured nerves. J. Cell Biol. 210, 153–168. 10.1083/jcb.201503019.

113. Guan, X.L., Cestra, G., Shui, G., Kuhrs, A., Schittenhelm, R.B., Hafen, E., van der Goot, F.G., Robinett, C.C., Gatti, M., Gonzalez-Gaitan, M., et al. (2013). Biochemical membrane lipidomics during Drosophila development. Dev. Cell 24, 98–111. 10.1016/j.devcel.2012.11.012.

114. Garbay, B., Heape, A.M., Sargueil, F., and Cassagne, C. (2000). Myelin synthesis in the peripheral nervous system. Prog. Neurobiol. 61, 267–304. 10.1016/s0301-0082(99)00049-0.

115. Svennerholm, L., and Vanier, M.T. (1978). Lipid and Fatty Acid Composition of Human Cerebral Myelin during Development. In Myelination and Demyelination, J. Palo, ed. (Springer US), pp. 27–41. 10.1007/978-1-4684-2514-7_3.

116. Svennerholm, L., Vanier, M.T., and Jungbjer, B. (1978). Changes in fatty acid composition of human brain myelin lipids during maturation. J. Neurochem. 30, 1383– 1390. 10.1111/j.1471-4159.1978.tb10470.x.

117. Zhu, Y., Cho, K., Lacin, H., Zhu, Y., DiPaola, J.T., Wilson, B.A., Patti, G.J., and Skeath, J.B. (2024). Loss of dihydroceramide desaturase drives neurodegeneration by disrupting endoplasmic reticulum and lipid droplet homeostasis in glial cells. eLife 13. 10.7554/eLife.99344.1.

118. Kilkus, J.P., Goswami, R., Dawson, S.A., Testai, F.D., Berdyshev, E.V., Han, X., and Dawson, G. Differential regulation of sphingomyelin synthesis and catabolism in oligodendrocytes and neurons.

119. McInnis, J.J., Sood, D., Guo, L., Dufault, M.R., Garcia, M., Passaro, R., Gao, G., Zhang, B., and Dodge, J.C. (2024). Unravelling neuronal and glial differences in ceramide composition, synthesis, and sensitivity to toxicity. Commun. Biol. 7, 1–17. 10.1038/s42003-024-07231-0.

120. Fernandes, V.M., Chen, Z., Rossi, A.M., Zipfel, J., and Desplan, C. (2017). Glia relay differentiation cues to coordinate neuronal development in Drosophila. Science 357, 886–891. 10.1126/science.aan3174.

121. Sasamura, T., Matsuno, K., and Fortini, M.E. (2013). Disruption of Drosophila melanogaster Lipid Metabolism Genes Causes Tissue Overgrowth Associated with Altered Developmental Signaling. PLoS Genet. 9, e1003917. 10.1371/journal.pgen.1003917.

122. Chen, J., Stork, T., Kang, Y., Nardone, K.A.M., Auer, F., Farrell, R.J., Jay, T.R., Heo, D., Sheehan, A., Paton, C., et al. (2024). Astrocyte growth is driven by the Tre1/S1pr1 phospholipid-binding G protein-coupled receptor. Neuron 112, 93–112.e10. 10.1016/j.neuron.2023.11.008.

123. Kang, Y., Jefferson, A., Sheehan, A., Torre, R.D.L., Jay, T., Chiao, L., Hulegaard, A., Corty, M., Baconguis, I., Zhou, Z., et al. (2023). Tweek-dependent formation of ER-PM contact sites enables astrocyte phagocytic function and remodeling of neurons. Preprint at bioRxiv, 10.1101/2023.11.06.565932 https://doi.org/10.1101/2023.11.06.565932.

124. Yin, J., Spillman, E., Cheng, E.S., Short, J., Chen, Y., Lei, J., Gibbs, M., Rosenthal, J.S., Sheng, C., Chen, Y.X., et al. (2021). Brain-specific lipoprotein receptors interact with astrocyte derived apolipoprotein and mediate neuron-glia lipid shuttling. Nat. Commun. 12, 2408. 10.1038/s41467-021-22751-7.

125. Hjelmqvist, L., Tuson, M., Marfany, G., Herrero, E., Balcells, S., and Gonzàlez-Duarte, R. (2002). ORMDL proteins are a conserved new family of endoplasmic reticulum membrane proteins. Genome Biol. 3, research0027.1. 10.1186/gb-2002-3-6-research0027.

126. Li, S., Xie, T., Liu, P., Wang, L., and Gong, X. (2021). Structural insights into the assembly and substrate selectivity of human SPT–ORMDL3 complex. Nat. Struct. Mol. Biol. 28, 249–257. 10.1038/s41594-020-00553-7.

127. Breslow, D.K., Collins, S.R., Bodenmiller, B., Aebersold, R., Simons, K., Shevchenko, A., Ejsing, C.S., and Weissman, J.S. (2010). Orm family proteins mediate sphingolipid homeostasis. Nature 463, 1048–1053. 10.1038/nature08787.

128. Clarke, B.A., Majumder, S., Zhu, H., Lee, Y.T., Kono, M., Li, C., Khanna, C., Blain, H., Schwartz, R., Huso, V.L., et al. (2019). The Ormdl genes regulate the sphingolipid synthesis pathway to ensure proper myelination and neurologic function in mice. eLife 8, e51067. 10.7554/eLife.51067.

129. Hagen-Euteneuer, N., Lütjohann, D., Park, H., Merrill, A.H., and Echten-Deckert, G. van (2012). Sphingosine 1-Phosphate (S1P) Lyase Deficiency Increases Sphingolipid Formation via Recycling at the Expense of de Novo Biosynthesis in Neurons *. J. Biol. Chem. 287, 9128–9136. 10.1074/jbc.M111.302380.

130. Tidhar, R., Ben-Dor, S., Wang, E., Kelly, S., Merrill, A.H., and Futerman, A.H. (2012). Acyl Chain Specificity of Ceramide Synthases Is Determined within a Region of 150 Residues in the Tram-Lag-CLN8 (TLC) Domain *. J. Biol. Chem. 287, 3197–3206. 10.1074/jbc.M111.280271.

131. Ohno, Y., Suto, S., Yamanaka, M., Mizutani, Y., Mitsutake, S., Igarashi, Y., Sassa, T., and Kihara, A. (2010). ELOVL1 production of C24 acyl-CoAs is linked to C24 sphingolipid synthesis. Proc. Natl. Acad. Sci. 107, 18439–18444. 10.1073/pnas.1005572107.

132. Bauer, R., Voelzmann, A., Breiden, B., Schepers, U., Farwanah, H., Hahn, I., Eckardt, F., Sandhoff, K., and Hoch, M. (2009). Schlank, a member of the ceramide synthase family controls growth and body fat in Drosophila. EMBO J. 28, 3706–3716. 10.1038/emboj.2009.305.

133. Rey, S., Ohm, H., Moschref, F., Zeuschner, D., Praetz, M., and Klämbt, C. (2023). Glial-dependent clustering of voltage-gated ion channels in Drosophila precedes myelin formation. eLife 12, e85752. 10.7554/eLife.85752.

134. Yunis, E.J., and Lee, R.E. (1970). Tubules of Globoid Leukodystrophy: A Right-Handed Helix. Science 169, 64–66. 10.1126/science.169.3940.64.

135. Inaba, T., Murate, M., Tomishige, N., Lee, Y.-F., Hullin-Matsuda, F., Pollet, B., Humbert, N., Mély, Y., Sako, Y., Greimel, P., et al. (2019). Formation of tubules and helical ribbons by ceramide phosphoethanolamine-containing membranes. Sci. Rep. 9, 5812. 10.1038/s41598-019-42247-1.

136. Murate, M., Tomishige, N., and Kobayashi, T. (2020). Wrapping axons in mammals and Drosophila: Different lipids, same principle. Biochimie 178, 39–48. 10.1016/j.biochi.2020.08.002.

137. Lorent, J., Levental, K., Ganesan, L., Rivera-Longsworth, G., Sezgin, E., Doktorova, M., Lyman, E., and Levental, I. (2020). Plasma membranes are asymmetric in lipid unsaturation, packing, and protein shape. Nat. Chem. Biol. 16, 644–652. 10.1038/s41589-020-0529-6.

138. Levental, I., Levental, K.R., and Heberle, F.A. (2020). Lipid Rafts: Controversies Resolved, Mysteries Remain. Trends Cell Biol. 30, 341–353. 10.1016/j.tcb.2020.01.009.

139. Kinoshita, M., Suzuki, K.G.N., Matsumori, N., Takada, M., Ano, H., Morigaki, K., Abe, M., Makino, A., Kobayashi, T., Hirosawa, K.M., et al. (2017). Raft-based sphingomyelin interactions revealed by new fluorescent sphingomyelin analogs. J. Cell Biol. 216, 1183– 1204. 10.1083/jcb.201607086.

140. Endapally, S., Frias, D., Grzemska, M., Gay, A., Tomchick, D.R., and Radhakrishnan, A. (2019). Molecular Discrimination between Two Conformations of Sphingomyelin in Plasma Membranes. Cell 176, 1040–1053.e17. 10.1016/j.cell.2018.12.042.

141. Róg, T., Orłowski, A., Llorente, A., Skotland, T., Sylvänne, T., Kauhanen, D., Ekroos, K., Sandvig, K., and Vattulainen, I. (2016). Interdigitation of long-chain sphingomyelin induces coupling of membrane leaflets in a cholesterol dependent manner. Biochim. Biophys. Acta 1858, 281–288. 10.1016/j.bbamem.2015.12.003.

142. Ziegler, A.B., Wesselmann, C., Beckschäfer, K., Wulf, A.-L., Dhiman, N., Soba, P., Thiele, C., Bauer, R., and Tavosanis, G. (2024). Lipid Disbalance Affects Neuronal Dendrite Growth and Maintenance in a Human Ceramide Synthase Disease Model. Preprint at bioRxiv, 10.1101/2024.10.31.621235 https://doi.org/10.1101/2024.10.31.621235.

143. Haney, M.S., Bohlen, C.J., Morgens, D.W., Ousey, J.A., Barkal, A.A., Tsui, C.K., Ego, B.K., Levin, R., Kamber, R.A., Collins, H., et al. (2018). Identification of phagocytosis regulators using magnetic genome-wide CRISPR screens. Nat. Genet. 50, 1716–1727. 10.1038/s41588-018-0254-1.

144. Zirin, J., Hu, Y., Liu, L., Yang-Zhou, D., Colbeth, R., Yan, D., Ewen-Campen, B., Tao, R., Vogt, E., VanNest, S., et al. (2020). Large-Scale Transgenic Drosophila Resource Collections for Loss- and Gain-of-Function Studies. Genetics 214, 755–767. 10.1534/genetics.119.302964.

145. Biswas, R., Stein, D., and Stanley, E.R. (2006). Drosophila Dok is required for embryonic dorsal closure. Development 133, 217–227. 10.1242/dev.02198.

146. Coutinho-Budd, J.C., Sheehan, A.E., and Freeman, M.R. (2017). The secreted neurotrophin Spätzle 3 promotes glial morphogenesis and supports neuronal survival and function. Genes Dev. 31, 2023–2038. 10.1101/gad.305888.117.

147. Öztürk-Çolak, A., Marygold, S.J., Antonazzo, G., Attrill, H., Goutte-Gattat, D., Jenkins, V.K., Matthews, B.B., Millburn, G., dos Santos, G., Tabone, C.J., et al. (2024). FlyBase: updates to the Drosophila genes and genomes database. Genetics 227, iyad211. 10.1093/genetics/iyad211.

148. Jenkins, V.K., Larkin, A., and Thurmond, J. (2022). Using FlyBase: A Database of Drosophila Genes and Genetics. In Drosophila: Methods and Protocols, C. Dahmann, ed. (Springer US), pp. 1–34. 10.1007/978-1-0716-2541-5_1.

149. Wagh, D.A., Rasse, T.M., Asan, E., Hofbauer, A., Schwenkert, I., Dürrbeck, H., Buchner, S., Dabauvalle, M.-C., Schmidt, M., Qin, G., et al. (2006). Bruchpilot, a protein with homology to ELKS/CAST, is required for structural integrity and function of synaptic active zones in Drosophila. Neuron 49, 833–844. 10.1016/j.neuron.2006.02.008.

150. Musashe, D.T., Purice, M.D., Speese, S.D., Doherty, J., and Logan, M.A. (2016). Insulin-like Signaling Promotes Glial Phagocytic Clearance of Degenerating Axons through Regulation of Draper. Cell Rep. 16, 1838–1850. 10.1016/j.celrep.2016.07.022.

151. Riedel, F., Gillingham, A.K., Rosa-Ferreira, C., Galindo, A., and Munro, S. (2016). An antibody toolkit for the study of membrane traffic in Drosophila melanogaster. Biol. Open 5, 987–992. 10.1242/bio.018937.

152. Klagges, B.R., Heimbeck, G., Godenschwege, T.A., Hofbauer, A., Pflugfelder, G.O., Reifegerste, R., Reisch, D., Schaupp, M., Buchner, S., and Buchner, E. (1996). Invertebrate synapsins: a single gene codes for several isoforms in Drosophila. J. Neurosci. Ok. J. Soc. Neurosci. 16, 3154–3165. 10.1523/JNEUROSCI.16-10-03154.1996.

153. Choi, S.W., Guan, W., and Chung, K. (2021). Basic principles of hydrogel-based tissue transformation technologies and their applications. Cell 184, 4115–4136. 10.1016/j.cell.2021.07.009.

154. Beachum, A.N., Salazar, G., Nachbar, A., Krause, K., Klose, H., Meyer, K., Maserejian, A., Ross, G., Boyd, H., Weigel, T., et al. (2024). Glia multitask to compensate for neighboring glial cell dysfunction. bioRxiv, 2024.09.06.611719. 10.1101/2024.09.06.611719.

